# Inflammation and matrix metalloproteinase 9 (Mmp-9) regulate photoreceptor regeneration in adult zebrafish

**DOI:** 10.1101/518365

**Authors:** Nicholas J. Silva, Mikiko Nagashima, Jingling Li, Laura Kakuk-Atkins, Milad Ashrafzadeh, David R. Hyde, Peter F. Hitchcock

## Abstract

Brain injury activates complex inflammatory signals in dying neurons, surviving neurons, and glia. Here, we establish that inflammation regulates the regeneration of photoreceptors in the zebrafish retina and determine the cellular expression and function of the inflammatory protease, matrix metalloproteinase 9 (Mmp-9), during this regenerative neurogenesis. Animals of either sex were used in this study. Following photoreceptor ablation, anti-inflammatory treatment suppresses the number of injury-induced progenitors and regenerated photoreceptors. Upon photoreceptor injury, *mmp-9* is induced in Müller glia, the intrinsic retinal stem cell, and Müller glia-derived photoreceptor progenitors. Deleting *mmp-9* results in over production of injury-induced progenitors and regenerated photoreceptors, but over time the absence of Mmp-9 compromises the maturation and survival of the regenerated cones. Anti-inflammatory treatment in mutants rescues the defects in cone maturation and survival. These data provide a link between injury-induced inflammation in the vertebrate CNS, Mmp-9 function during photoreceptor regeneration and the requirement of Mmp-9 for the survival of regenerated cones.

**Significance Statement:** The innate immune system is activated by neuronal death, and recent studies demonstrate that in zebrafish neuroinflammation is required for neuronal regeneration. The roles of inflammatory cytokines are being investigated, however, the function of the inflammatory protease, matrix metalloprotease Mmp-9, in neuronal regeneration is unknown. We show herein that in adult zebrafish retinal inflammation governs the proliferative phase of the stem cell-based regeneration of rod and cone photoreceptors and determine the specific roles for Mmp-9 in photoreceptor regeneration. This study provides the first mechanistic insights into the potential role of Mmp-9 in retinal regeneration and serves to link neuroinflammation, stem cell-based regeneration of photoreceptors and human photoreceptor disease.

## Introduction

Inflammation modulates immune and nonimmune functions during tissue development, repair, and diseases (Deverman and Patterson, 2009; Ekdahl et al., 2009). Acute inflammation involves the secretion of proinflammatory cytokines and chemokines that recruit immune cells to the damaged tissue (Liddiard et al., 2011; Nathan, 2002). In the central nervous system of adult mammals, acute inflammation can activate signaling cascades in stem and progenitor cells that stimulate proliferation and promote neurogenesis (Borsini et al., 2015; Ekdahl et al., 2009; Kizil et al., 2015; Kyritsis et al., 2014, 2012). Chronic inflammation, however, is detrimental, and results in secondary damage to neurons and, in severe cases, neurodegenerative disease (Amor et al., 2010). Proper regulation of the inflammatory cascades in the central nervous system is critical both to maintain tissue homeostasis and successfully repair brain injuries.

Unlike mammals, which possess a limited capacity for tissue regeneration, zebrafish have an astonishing ability to regenerate tissues and organs, such as fins, heart, brain, and retina (Gemberling et al., 2013; Kizil et al., 2012; Lenkowski and Raymond, 2014). Studies using zebrafish have identified cellular and molecular mechanisms which may unlock the regenerative potential of mammalian tissues (Ueki et al., 2015). Such mechanisms include inflammation. In zebrafish acute inflammation is both necessary and sufficient to induce neuronal regeneration (Kyritsis et al., 2012; White et al., 2017; Caldwell et al., 2019).

In the zerbrafish retina, Müller glia serve as intrinsic stem cells that underlie the ability to regenerate retinal neurons (Bernardos et al., 2007; Fausett and Goldman, 2006). Neuronal injury and death stimulate Müller glia to initiate a transient gliotic response, followed by partial dedifferentiation and entry into the cell cycle (Raymond et al., 2006; Thomas et al., 2016). Müller glia then undergo a single asymmetric division to produce multipotent progenitors, which proliferate rapidly, migrate to areas of cell death and differentiate to replace the ablated neurons (Fimbel et al., 2007; Sherpa et al., 2008; Goldman, 2014; Gorsuch and Hyde, 2014; Lenkowski and Raymond, 2014). In the zebrafish retina, paracrine and autocrine signaling that engage Müller glia plays an essential role facilitating this regenerative neurogenesis (Nelson et al., 2013; Zhao et al., 2014). Following a photolytic lesion, dying photoreceptors secrete Tnf-α, to which Müller glia respond by partial dedifferentiation, synthesis of Tnf–α and entry into the cell cycle (Nelson et al., 2013). Mechanical lesions induce the expression of *leptin* and Il-6 family cytokines in Müller glia and Müller glia-derived progenitors and is required for injury-induced proliferation (Zhao et al., 2014).

Matrix metalloproteinase 9 (Mmp-9) is a secreted protease that plays a prominent role in tissue development and homeostasis by acting on extracellular molecules, including adhesion molecules, growth factors and cytokines (Bonnans et al., 2014; Le et al., 2007; Masure et al., 1991; Parks et al., 2004; Vandooren et al., 2013b, 2014), which, in turn, can regulate proliferation, cellular migration and cellular differentiation. It is well established that inflammatory cytokines can induce the expression of MMPs, including MMP-9 (Vecil et al., 2000; Shubayev et al., 2006; Nagase, et al., 2006; Vandooren, et al., 2013b). MMP-9 was first purified from neutrophils and monocytes following stimulation by IL-8 and IL-1β, implicating its role as an inflammatory protease (Masure et al. 1991; Opdenakker, et al., 1991). In adult tissues, Mmp-9 is dramatically induced following various types of injuries (Vandooren et al., 2013b, 2014; Xu et al., 2018). The complexities of Mmp-9 function suggest its role during regeneration is likely tissue, substrate and injury context dependent (Hindi et al., 2013; Vandooren et al., 2014; Ando et al., 2017; Xue et al., 2017). The cellular expression and function of Mmp-9 during photoreceptor regeneration in the zebrafish are unknown.

Here, we determine the general role of inflammation and the cellular expression and function of the matrix metalloproteinase, Mmp-9, during photoreceptor regeneration. We establish that in the adult retina inflammation governs the proliferation of Müller glia-derived progenitors. We show that *mmp-9* is expressed in Müller glia, as they prepare to enter the cell cycle, and Müller glia-derived progenitors. Using a loss-of-function mutant, we determine that Mmp-9 negatively regulates the proliferation of Müller glia-derived progenitors. Finally, we demonstrate that following photoreceptor regeneration, Mmp-9 is required for the maturation and survival of regenerated cone photoreceptors.

## MATERIALS AND METHODS

### Animals

Wild-type, AB strain zebrafish (*Danio rerio*; ZIRC, University of Oregon, Eugene, OR) were propagated, maintained, and housed in recirculating habitats at 28.5°C and on a 14/10-h light/dark cycle. Embryos were collected after natural spawns, incubated at 28.5°C and staged by hours post fertilization (hpf). Adults were of either sex and used between 6 and12 months of age. The transgenic reporter line, *Tg*[*gfap:EGFP*]*^mi2002^*, was used to identify Müller glia in retinal sections (Bernardos and Raymond, 2006). All experimental protocols were approved by the University of Michigan’s Institutional Animal Care and Use Committee (IACUC).

### Light Lesions

To kill photoreceptors, fish were dark adapted, then exposed to high intensity light (ca. 100,000 lux) for 30 minutes, followed by exposure to constant light (ca. 30.000 lux) for 72 hours (Taylor et al., 2015). After 72 hours fish were returned to the recirculating habitats and normal light/dark cycle.

### Plastic sections

Eye cups were fixed overnight at 4°C in 2% PFA/2% glutaraldehyde in 0.1M phosphate buffer, then infiltrated and embedded in glycol methacrylate plastic resin (JB-4; Polysciences, Inc., Warrington, PA). Sections were cut at 4-μm, mounted on glass slides and stained with 0.13% methylene blue and 0.13% basic fuchsin (Lee’s stain).

### Quantitative Real-Time PCR

Total RNA was extracted from dissected whole retinas using TRIzol (Invitrogen, Carlsbad, CA). Reverse transcription to cDNA was performed using 1 µg RNA (Qiagen QuantiTect Reverse Transcription kit; Venlo, Netherlands), and each qPCR reaction was performed with 6 ng cDNA and Bio-Rad IQ SYBR Green Supermix (Bio-Rad CFX384 Touch Real Time PCR Detection System; Hercules, CA). Three independent biological replicates were collected at each time point, and each replicate contained six homogenized retinas from three fish (18 retinas/time point). Each biological replicate was run in triplicate. For each time point, average mRNA levels were represented as Log2 fold change, calculated using the DDC_T_ method and normalized to the housekeeping gene, *gpia*. Primers used for qPCR analyses are listed in Table 1.

**Table 1:**
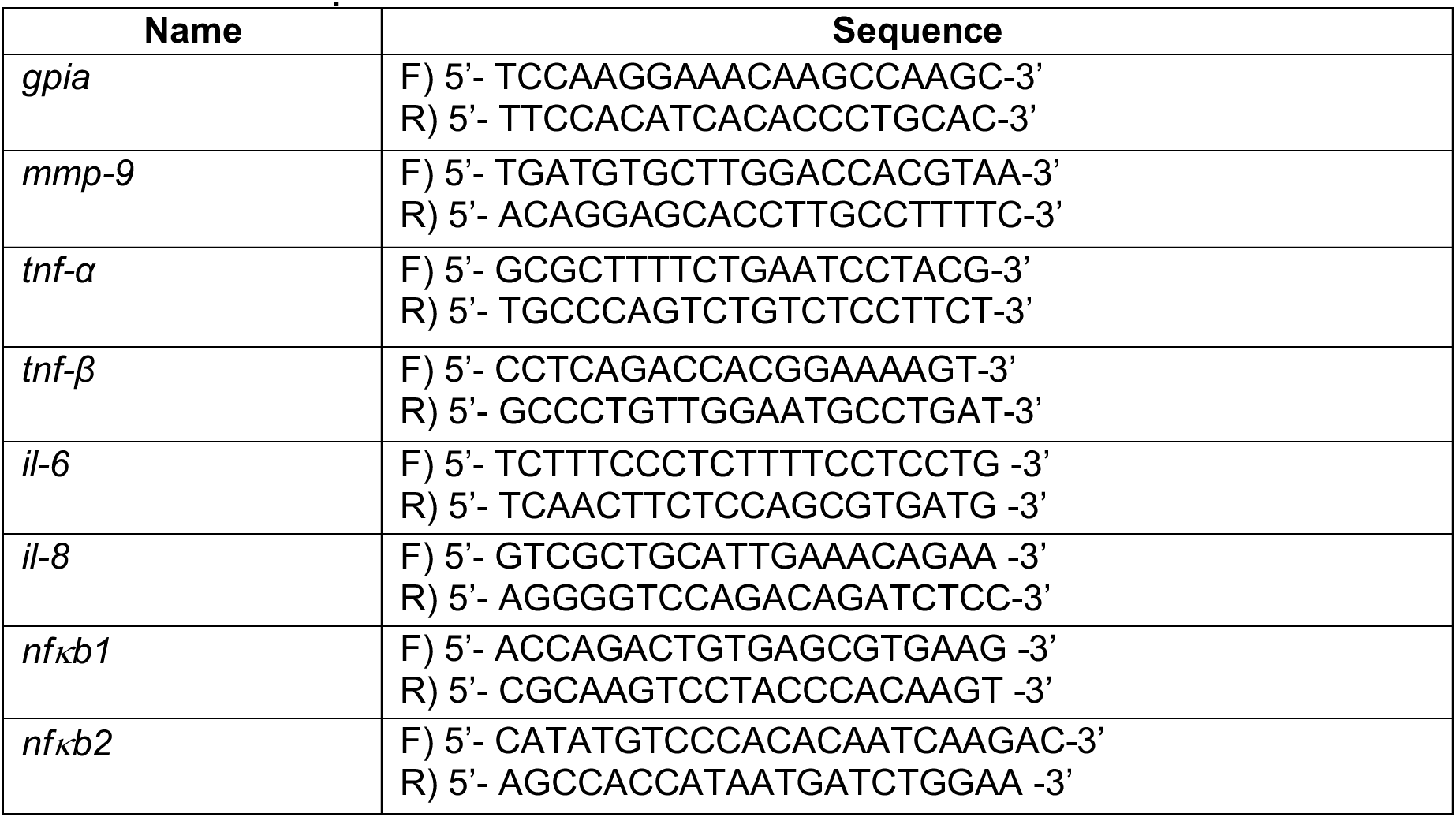
Primer Sequences

### Dexamethasone treatment

A previously described protocol was used that efficiently suppresses the immune system in zebrafish (Kyritsis et al., 2012; see also Bollaerts et al., 2019). Briefly, fish were housed in system water with 15 mg/L dexamethasone (Sigma-Aldrich, Corp, D1756) diluted in 0.1% MetOH. Dexamethasone-containing and control solutions were changed daily, and fish were fed brine shrimp every other day.

### Immunohistochemistry

Larvae or dissected eyecups were fixed overnight at 4°C in 0.1M phosphate buffered 4% paraformaldehyde, cryoprotected with 20% sucrose, and embedded in optical cutting temperature (OCT) medium (Sakura Finetek USA, Torrance, CA). Immunohistochemistry (IHC) was performed as previously described (Taylor et al., 2015). Briefly, 10-μm-thick sections were collected such that the optic nerve head was included, and sections were mounted onto glass slides. Sections were washed in phosphate buffer saline with 0.5% Triton-x (PBST) and incubated with 20% heat-inactivated normal sheep serum in PBST for 2 hours (NSS; Sigma-Aldrich, Corp.). Primary antibodies were applied overnight at 4°C. Sections were then washed with PBST and incubated in secondary antibodies for 1 hour at room temperature. Prior to IHC for BrdU, sections were immersed in 100°C sodium citrate buffer (10 mM sodium citrate, 0.05% Tween 20, pH 6.0) for 30 minutes and cooled at room temperature for 20 minutes. Sections were then subjected to IHC as described above.

Immunohistochemistry performed on whole retinas was conducted as described previously (Nagashima et al., 2017). Prior to IHC, retinas were dissected from dark-adapted animals, flattened with radial relaxing cuts, fixed at 4°C overnight in 4% paraformaldehyde in 0.1M phosphate buffer with 5% sucrose. Retinas were then placed in boiling sodium citrate buffer for 5 min and allowed to cool for 5 min. Retinas were incubated for 2 hours in 10% normal goat serum, 1% Tween, 1% Triton X-100, 1% DMSO, in phosphate buffered saline with 0.1% sodium azide. Primary and secondary antibody were diluted in 2% normal goat serum, 1% Tween, 1% Triton X-100, 1% DMSO, in phosphate buffered saline with 0.1% sodium azide and incubations were performed overnight at room temperature. After washing in phosphate buffered saline with 1% Tween, 1% Triton X-100, 1% DMSO, retinas were mounted on glass slides, photoreceptor side down, in ProLong Gold anti-fade reagent (Life Technologies Corp., Eugene, OR). Antibodies used in this study and their concentrations are listed in Table 2.

**Table 2:** Antibody List

| **Primary Antibodies** | **Company** | **Dilution** |
| --- | --- | --- |
| Mouse monoclonal anti-GFP | Milipore<br>MAB3580 | 1:200 |
| Mouse monoclonal anti-actin | Calbiochem | 1:1000 |
| Mouse Monoclonal anti-Zn5 | ZIRC; zfin.org/ZDB-ATB-081002-19 | 1:200 |
| Mouse Monoclonal anti-Zpr1 (anti-Arrestin-3) | ZIRC; zfin.org/ZDB-ATB-081002-43 | 1:200 |
| Mouse Monoclonal anti-Zpr-3 | ZIRC; zfin.org/ZDB-ATB-081002-45 | 1:200 |
| Mouse anti-HPC1 | ZIRC; zfin.org/ZDB-ATB 130225-1 | 1:200 |
| Mouse anti-Zrf1 | ZIRC; zfin.org/ZDB-ATB-081002-46 | 1:1000 |
| Mouse Monoclonal ZO-1A-12 | Innovative Research Cat# 33-9100, RRID:AB_87181 | 1:200 |
| Mouse anti-BrdU | BD Biosciences Ca# 347580 | 1:100 |
| Rat anti-BrdU | Abcam Cat# ab6326, RRID:AB_305426 | 1:200 |
| Polyclonal anti-GNAT2 | MBL Ca# PM075 | 1:1000 |
| Mouse Monoclonal anti-1D1 (ZF Rhodopsin) | Hyatt et al., 1996<br>Gift from Dr. Jim Fadool (FSU) | 1:1000 |
| Mouse Monoclonal anti-4C4 (zebrafish microglia) | Raymond et al., 2006<br>Hitchcock Lab | 1:200 |
| **Secondary Antibodies** |  |  |
| Alexa Fluor goat anti-mouse 488, 555, or 647 | Invitrogen | 1:500 |
| Alexa Fluor goat anti-rabbit 488, 555, or 647 | Invitrogen | 1:500 |
| Alexa Fluor goat anti-rat 488, 555, or 647 | Invitrogen | 1:500 |

### Labeling dividing cells

For embryos, dividing cells were labeled with EdU by soaking embryos for 20 min in 1.5 mM EdU dissolved in embryo rearing solution containing 15% DMSO. EdU was visualized using the Click-iT™ EdU imaging kit (Thermo Fisher Scientific, Waltham, MA) with Alex Fluor 555 according to the manufacturer’s protocol. In adults, dividing cells were labeled with BrdU by housing animals in system water containing 5 mM BrdU for 24 hours (Gramage et al., 2015).

### *In situ* Hybridization

*In situ* hybridizations were performed as previously described (Luo et al., 2012). Digoxigenin (DIG)-labeled riboprobes for *rhodopsin* and *pde6c* were generated from full-length cDNA clones (Luo et al., 2012; Ochocinska and Hitchcock, 2007), whereas riboprobe for *mmp-9* was generated by PCR amplification using primers containing the T3 or T7 promoter sequences (David and Wedlich, 2001). *mmp-9* (F): 5′<u>TAATACGACTCACTATAGGG</u>GATTCTTCTACTTCTGCCGGG 3′ *mmp-9* (R): 5′<u>AATTAACCCTCACTAAAGGG</u>CTTAATAAATTTGTAAACAAG 3′.

Briefly, 10-μm-thick sections were hybridized with riboprobes at 55°C, incubated with an alkaline-phosphatase-conjugated anti-DIG antibody and visualized using Fast Red TR/Naphthol AS-MX (SIGMA*FAST*) as the enzymatic substrate. When *in situ* hybridizations were combined with BrdU IHC, sections were removed from the fast red solutions, rinsed and post-fixed in buffered 4% paraformaldehyde for 10 minutes then processed for BrdU IHC as described above.

### Production of custom antibodies

Antibodies specific to zebrafish Mmp-9 were generated by Pocono Rabbit Farm & Laboratory (PRF&L, Canadensis, PA) as previously described (Calinescu et al., 2009). A 24 amino acid C-terminal peptide was used as the immunogen, C<u>DIDGIQYLYGPRTGPEPTAPQPR</u>; NCBI: AY151254. Polyclonal antibodies were affinity purified and confirmed by ELISA (data not shown). Western blots performed with retinal proteins using pre- and post-immune sera confirmed the post-immune serum detected a 76 kDa band, the predicted size for Mmp-9. Western blot analysis and known cleavage sites on pro-Mmp9 suggest these antibodies recognize both the pro- and active forms of Mmp-9, which can be distinguished by slight differences in their molecular weights (Vandooren et al., 2013a; see Results).

### Western Blot Analysis

Protein samples were obtained from whole retinas homogenized in RIPA lysis buffer (ThermoFisher Scienific) containing protease and phosphatase inhibitor cocktail (5872S; Cell Signaling Technology, Danvers, MA, USA). Proteins were separated in a 12% Mini-PROTEAN TGX Precast gel (BioRad) and transferred to a polyvinylidene difluoride (PVDF) membrane (GenHunter Corp., Nashville, TN). To block non-specific staining, membranes were incubated in 5% nonfat dry milk in Tris buffered saline containing 0.3% Tween-20 (TBST) for 2 hours. Membranes were incubated with the antibodies-containing solution overnight at 4°C. Blots were then washed in TBST and incubated with horseradish peroxidase-conjugated secondary IgG (1:1000) for 1 hour at room temperature. Antibody-bound protein was visualized using Pierce ECL Western blotting substrate (32106; ThermoFisher Scienific, Waltham, MA). To visualize loading controls, blots were also stained with antibodies against actin. Images were captured using the Azure C500 (Azure Biosystems, Dublin, CA). Densitometry of protein bands was performed using ImageJ software (https://imagej.nih.gov/ij/).

### Zymography

Protein samples were prepared as described for the Western blot analyses. Proteins were separated on 10% Zymogram Plus (Gelatin) Protein Gels, (ThermoFisher Scientific). Following electrophoresis, proteins were renatured in 1X renaturing buffer (ThermoFisher Scientific) for 30 mins at room temperature, then incubated in 1X developing buffer (ThermoFisher Scientific) overnight at 37°. Gels were rinsed in deionized water and stained with SimplyBlue SafeStain (ThermoFisher Scientific). Gels were imaged with long-wave ultraviolet light using the Azure C500 (Azure Biosystems; Dublin, CA). Active recombinant human MMP-9 (Calbiochem, San Diego, CA) was used as a positive control. Densitometry of the digested bands was performed using ImageJ software.

### CRISPR-Cas9 mediated targeted mutation of Mmp-9

*mmp-9* mutants were generated according to previously described methods (Hwang et al., 2013). ZiFiT software (available in the public domain at zifit.partners.org) was used to identify a 19 bp sgRNA target sequence for *mmp-9* (GGCTGCTTCATGGCATCAA). Target sequence Oligo1-TTGATGCCATGAAGCAGCC and Oligo2-AAAC<u>TTGATGCCATGAAGCAG</u> were annealed and subcloned into the pT7 gRNA vector (Addgene ID: 46759). The pCS2 nCas9n vector (Addgene ID: 46929) was used for Cas9 synthesis. To produce RNAs, the MEGAshortscript T7 kit (Ambion, Foster City, CA: AM1354) and mirVana miRNA Isolation kit (Ambion: AM 1560) were used for the gRNA, whereas, mMesage mMACHINE SP6 kit (Ambion: AM1340) and RNeasy mini Kit from (Qiagen: 74104) were used for the Cas9 mRNA. Single cell-stage embryos were injected with a 2nL solution containing 100 pg/nl sgRNA and 150 pg/nl Cas9 mRNA. Founders (F0) were outcrossed to AB wild-type fish. Mutations were identified using screening primers (F: 5′-AAGTCTGCAACTACATATCAGC-3′, R: 5′-GTACACACTGTAGATGCTGATAAG-3′) that flanked the *mmp-9* sgRNA target site. Standard protocols for PCR used Platinum Taq-HF polymerase (Invitrogen) with genomic DNA as the template. The purified PCR product was subcloned into the pGEM-T Easy vector (Promega, Madison, WI) for Sanger sequencing. Mutant and wild-type sequences were aligned using pairwise blast (National Center for Biotechnology Information [NCBI], Bethesda, MD, USA). Premature stop codons were identified by comparing predicted amino acid sequences for wild-type and mutants using the ExPASy translate tool (available in the public domain at (www.expasy.org). F1 hybrids carrying a *mmp-9* mutation were in-crossed and homozygous F2 mutants were identified by a combination of Sanger sequencing and T7 endonuclease assays (New England Biolabs, Ipswich, MA) as previously described (www.crisprflydesign.org).

### Imaging

Fluorescence images of retinal sections were captured using the Leica TCS SP5 confocal microscope (Leica Microsystems, Wetzlar, Germany). Cell counts were conducted using z-stack images and analyzed with Imaris software (Bitplane, South Windsor, CT). Regenerated photoreceptors were identified by the colocalization of DAPI, BrdU, and the ISH markers *pde6c* (cones) or *rho* (rods).

### Cell Counts and area measurement

Cells were counted in both radial sections and retinal wholemounts. In radial sections, cells were counted in three nonadjacent sections in each retina. The number of cells per retina were averaged, and averages were computed for control and experimental groups. In wholemounts stained with an antibody against the tight junction protein, ZO-1 (Nagashima et al., 2017), cones were identified in optical sections taken at the outer limiting membrane as profiles with perimeters greater than 3.5 μm. Cells were counted in five separate regions in each retina, sampling a totaling 5625^2^μm per retina. The average number of cells in control and experimental groups were then averaged.

### Experimental Design and Statistical Analyses

The statistical significance in mRNA levels between unlesioned and lesioned retinas was calculated using an ANOVA (JMP 9.0; SAS Institute, Inc., Cary, NC). A *p*-value ≤ 0.05 was considered significant. The statistical significance in the number of cells between control and treated retinas was calculated using a student’s *t*-test (GraphPad Prism Software, La Jolla, CA). A *p*-value ≤ 0.05 was considered significant. Western and zymogram analyses consisted of three independent biological replicates, and each replicate contained six homogenized retinas from three fish (18 retinas per time point). Statistical significance between optical density values of bands in control and experimental groups was calculated using an ANOVA (JMP 9.0, SAS Institute, Inc., Cary, NC). A *p*-value ≤ 0.05 was considered significant.

## RESULTS

### The expression of inflammatory genes is induced by photoreceptor ablation

To characterize the inflammatory response during photoreceptor ablation and regeneration, qPCR was used to quantify the expression of inflammatory genes, *mmp-9, tnf*–*α*, *tnf-β, il-8, nfκb1*, and *nfκb2*, shown previously to regulate various forms of tissue regeneration in zebrafish (de Oliveira et al., 2013; LeBert et al., 2015; Nelson et al., 2013; Karra et al., 2015). The time course of gene expression follows closely the well-established time course for the injury response in Müller glia and the appearance of the Müller glia-derived progenitors and their subsequent differentiation into mature photoreceptors (Figure 1A; see (Gorsuch and Hyde, 2014)). Upregulation of gene expression can be detected by 8 hours post lesion (hpl), and the rising phase in expression corresponds to the period when Müller glia prepare to enter the cell cycle. Expression levels then slowly decline as the Müller glia-derived photoreceptor progenitors divide, migrate to the outer nuclear layer and differentiate into regenerated photoreceptors (168 hpl; Figure 1A). Interestingly, among all the transcripts examined, *mmp-9* shows the greatest change from the undamaged controls, peaking at 24-48 hpl. Further, by 168 hpl *mmp-9* levels are reduced to only about 70% of the peak value (ANOVA F-ratio = 22.23, p=.0001; Figure 1A). The pro-inflammatory cytokine, Tnf-α, has been posited to be responsible for initiating photoreceptor regeneration in zebrafish (Gorsuch and Hyde, 2014; Nelson et al., 2013), but at all time points in this assay, the levels of *tnf-α* expression were not significantly different from controls. Our assay for *tnf-α* failed to replicate previous results (Nelson et al., 2013), and we infer this is most likely a consequence of technical issues, perhaps reflecting differences between the qPCR assay used here and the protein analysis used by Nelson et al. (2013). Nonetheless, these data show that in zebrafish a photolytic lesion, which leads to photoreceptor death, induces rapid expression of characteristic inflammatory genes that follows a time course reflecting the well-described events that underlie photoreceptor regeneration. Further, the expression of *mmp-9* remains significantly elevated as photoreceptor differentiation commences.

**Figure 1:**
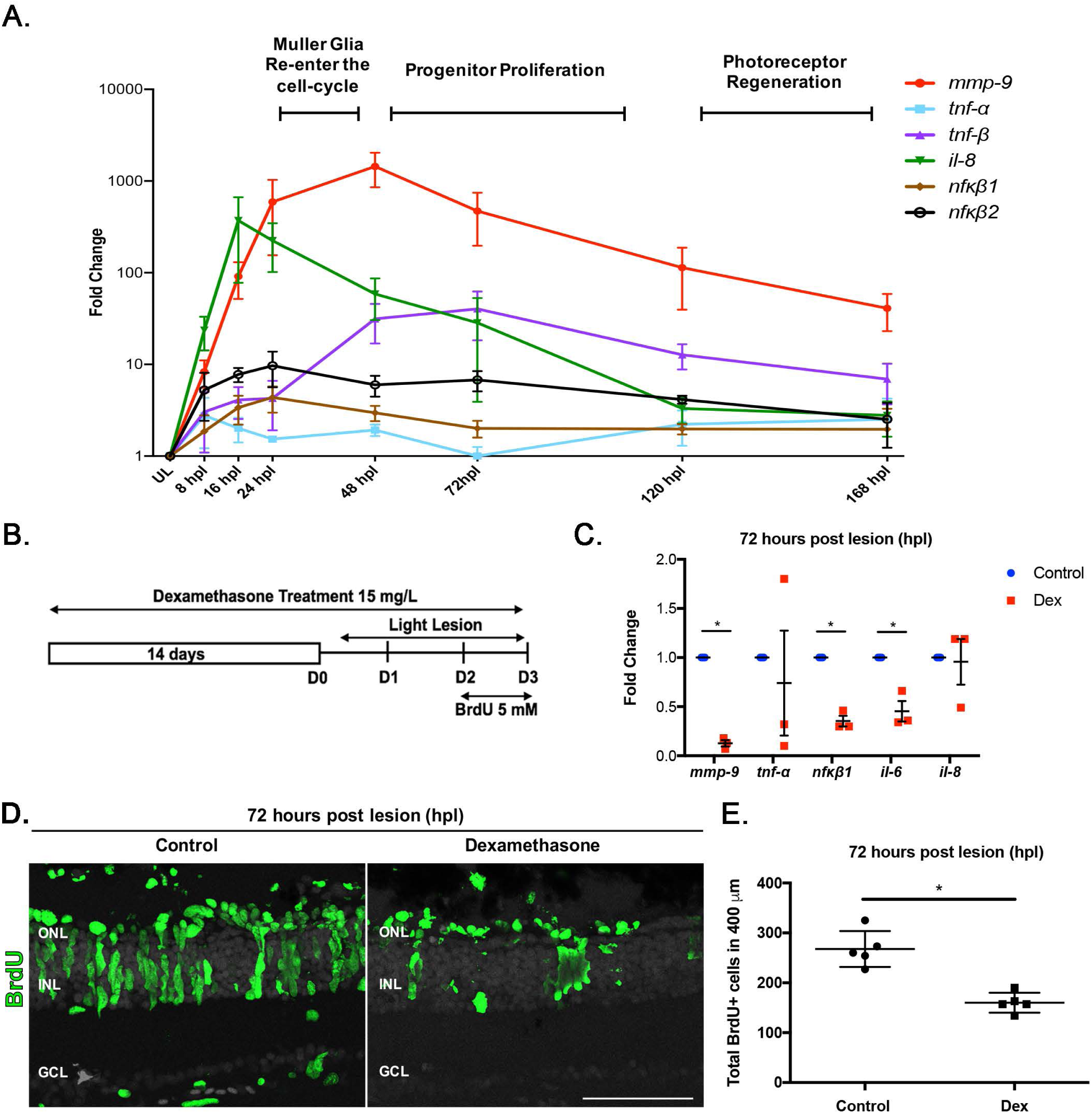
Inflammation is induced following photoreceptor ablation and regulates reactive-proliferation. **(A)** Time-course for the expression of inflammatory genes, from 8 to 168 hours post lesion (hpl). Unlesioned retinas served as controls. Expression levels are represented as fold change calculated using DD*C*_T_ method. **(B)** Experimental paradigm for anti-inflammatory treatment with Dex and BrdU labeling. **(C)** qRT-PCR for the inflammatory genes *mmp-9, tnf-a, nfkb1, il-8*, and *il-6* from control and Dex-treated retinas at 72 hpl. *p≤0.05. **(D)** BrdU immunostained cells (green) in controls (left) and Dex-treated animals (right) at 72 hpl. **(E)** Number of BrdU-labeled cells in controls (268 ± 36.1 cells; n=5) and Dex-treated animals (160.2 ± 20.02 cells; n=5) at 72 hpl; *p=0.0079. ONL-outer nuclear layer; INL-inner nuclear layer; GCL-ganglion cell layer. Scale bar equals 50***μ***m.

### Inflammation regulates injury-induced proliferation and photoreceptor regeneration

To determine if inflammation regulates aspects of photoreceptor regeneration, the glucocorticoid steroid, dexamethasone (Dex), was used to suppress the inflammatory response. A 14-day pre-lesion treatment with dexamethasone was chosen based on studies performed by Kyritsis et al. (2012; see also White et al., 2017; Caldwell et al., 2019; Bollaerts et al., 2019), and here the dose of dexamethasone, route of administration and sizes and ages of animals were matched with Kyritis et al. (2012). Whereas this lengthy treatment with dexamethasone may affect homeostatic function in all cells (Juszczak and Stankiewicz, 2018), this treatment in the absence of a lesion does not directly alter constitutive proliferation or neurogenesis (Kyritsis et al., 2012; White et al., 2017; Caldwell et al., 2019).

To validate the effectiveness of the Dex treatment, the expression levels of inflammatory genes were quantified in control and Dex-treated groups. At 72 hpl, the expression of *mmp-9*, *il*-*6*, and *nfκb1* were significantly decreased following dexamethasone treatment (p values = .0001, .0061, .0003, respectively; Figure 1C). Dex treatment did not suppress the expression of *tnf-α*, and there was marked variability in the qPCR results for *tnf-α*. The reasons for the failure of the Dex treatment to suppress *il-8* expression are unresolved. Further, Dex suppression of NFkB mediated gene transcription may directly or indirectly affect mmp9 expression and/or activity.

Assays of cell proliferation and photoreceptor regeneration were performed for control animals and animals treated with Dex. Dividing cells were labelled with BrdU between 24 and 48 hpl to mark progenitors that will give rise to regenerated photoreceptors (Figure 1B), and regenerated photoreceptors were identified as BrdU-labeled nuclei surrounded by *in situ* hybridization signal for either *rho* (rods) or *pde6c* (cones) at 7 dpl (Figure 2A). At 72 hpl, there was significantly fewer BrdU-labeled cells in Dex-treated retinas compared to controls (p=0.0079; Figure 1D, E). At 7 dpl, there were significantly fewer regenerated rod and cone photoreceptors in Dex-treated groups compared to controls (Figure 2B, C; p=0.0012). These results indicate that, consistent with data showing the requirement for inflammation in neuronal regeneration in the forebrain of zebrafish (Kyritsis et al., 2012), inflammatory mechanisms also promote photoreceptor regeneration in the retina.

**Figure 2:**
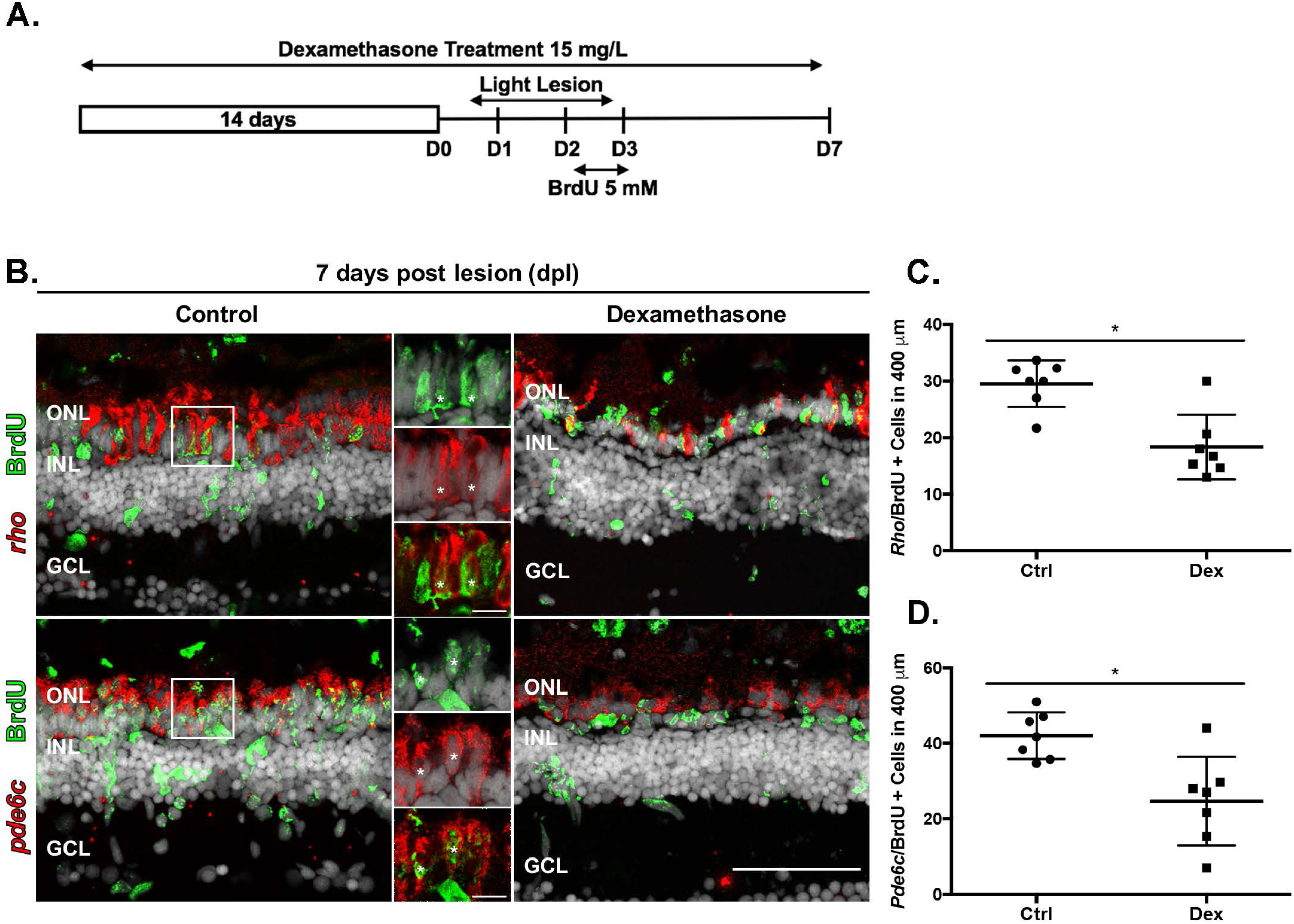
Anti-inflammatory treatment suppresses photoreceptor regeneration. **(A)** Experimental paradigm of anti-inflammatory treatment. **(B)** Double labeled, regenerated photoreceptors using *in situ* hybridization for rods (*rho) and cones (pde6c;* red signal) and BrdU (green). The high magnification insets show the colocalization of the two labels (asterisks). **(C)** Number of regenerated rod photoreceptors in control (29.52 ± 4.1 cells; n=7) and experimental retinas (1833 ± 5.71 cells; n=7); *p=0.0047. **(D)** Number of regenerated cone photoreceptors in control (42 ± 6.1 cells; n=7) and Dex-treated animals (25 ± 11.72 cells; n=7). *p=0.0012. ONL-outer nuclear layer; INL-inner nuclear layer; GCL-ganglion cell layer. Scale bar equals 50***μ***m.

### Müller glia and Müller glia-derived progenitors express *mmp-9*

MMP-9 belongs to a large family of Zn^2^+-dependent proteases that target a variety of substrates, such as extracellular matrix (ECM), growth factors, chemokines, and cytokines. Mmp-9 serves diverse functions among vertebrates (Pedersen et al. 2015; Page-McCaw, et al., 2007; Vandooren, 2013b; Iyer et al. 2012). Due to the varied roles played by *mmp-9* in injured/regenerating tissues (Vandooren et al., 2014), and its persistent expression throughout the time-course of photoreceptor regeneration (Figure 1A), the specific function of Mmp-9 was investigated further. As a first step, *in situ* hybridization was performed to identify the cellular patterns of expression of *mmp-9*. This was undertaken using the Müller glia reporter line, *Tg*[*gfap:EGFP*]*mi2002* (Bernardos and Raymond, 2006), and BrdU labeling 24 hrs prior to sacrifice. By *in situ* hybridization, *mmp-9* is not detectable in unlesioned retinas. In contrast, by 24 hpl, *mmp-9* is expressed exclusively in Müller glia, but at a time-point prior to their entry into S-phase of the cell cycle (Figure 3A). By 48 hpl, ∼97% of BrdU-labeled Müller glia express *mmp-9* (Figure 3A, B). At 72 hpl, the expression of *mmp-9* decreases (Figure 3A, C), and the cellular expression shifts to the Müller glia-derived progenitors, as evidenced by the absence of eGFP in BrdU-labeled cells that express *mmp-9* (Figure 3A). These data indicated that photoreceptor ablation induces the expression of *mmp-9* in Müller glia, and this expression persists in Müller glia-derived progenitors. Further, these results suggest that *mmp-9* identifies the subset of Müller glia that enter the cell cycle in response to a retinal injury.

**Figure 3:**
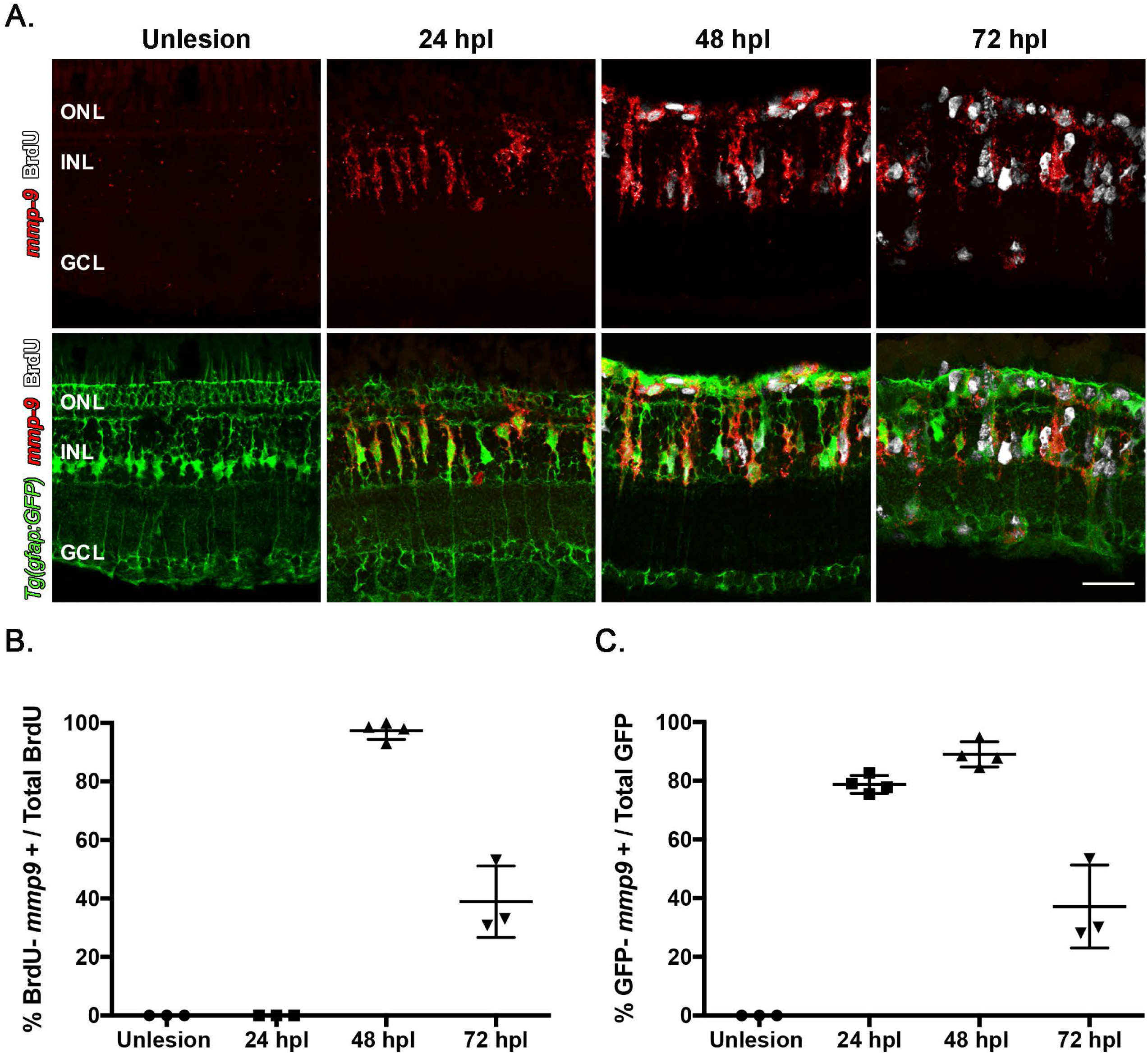
Following a photolytic lesion, Müller glia and Müller glia-derived progenitors express *mmp-9*. These experiments were performed using the transgenic line *Tg[gfap:EGFP]mi2002*, in which eGFP is expressed in Müller glia. **(A)** Triple labeling using *in situ* hybridization for *mmp-9* (red) and immunostaining for BrdU for dividing cells (white), combined with GFP immunostaining (green). **(B)** Number of BrdU+ cells that express *mmp-9* in unlesioned animals and lesioned animals at 24, 48, and 72 hpl. **(C)** Number of GFP+ Müller glia that also express *mmp-9* expression at 24, 48, and 72 hpl. ONL-outer nuclear layer; INL-inner nuclear layer; GCL-ganglion cell layer. Scale bar equals 25***μ***m.

### Mmp-9 protein is present and catalytically active following photoreceptor death

In tissues, Mmp-9 is synthesized and secreted in a pro-form, as a zymogen, then cleaved into an active protease, which can be distinguished by the slight difference in its molecular weight (Chattopadhyay and Shubayev, 2009; Vandooren et al., 2013a). Western blot analysis was used to characterize the induction of protein synthesis and cleavage, and zymogram assays were used to determine the time course of protein synthesis and catalytic activity for Mmp-9 during photoreceptor regeneration (Vandooren et al., 2013a, 2013b). By these assays, Mmp-9 is not detectable in unlesioned retinas, consistent with the *in situ* hybridization data (Figure 3A, 4A). At 16hpl, the first time point sampled, the un-cleaved, pro-form of Mmp-9 is detected as an upper band in Western blots (Figure 4A). Mmp-9 levels peak by 24hpl, and both forms of Mmp-9 are detected. Mmp-9 levels decrease between 24 and 72hpl, and during this interval the protein mostly appears in a slightly lower band, consistent with the active form of the protein. Mmp-9 levels are undetectable by Western blot analysis at 5dpl (ANOVA F-ratio = 8.377, p =.0013) (Figure 4 A, B).

**Figure 4:**
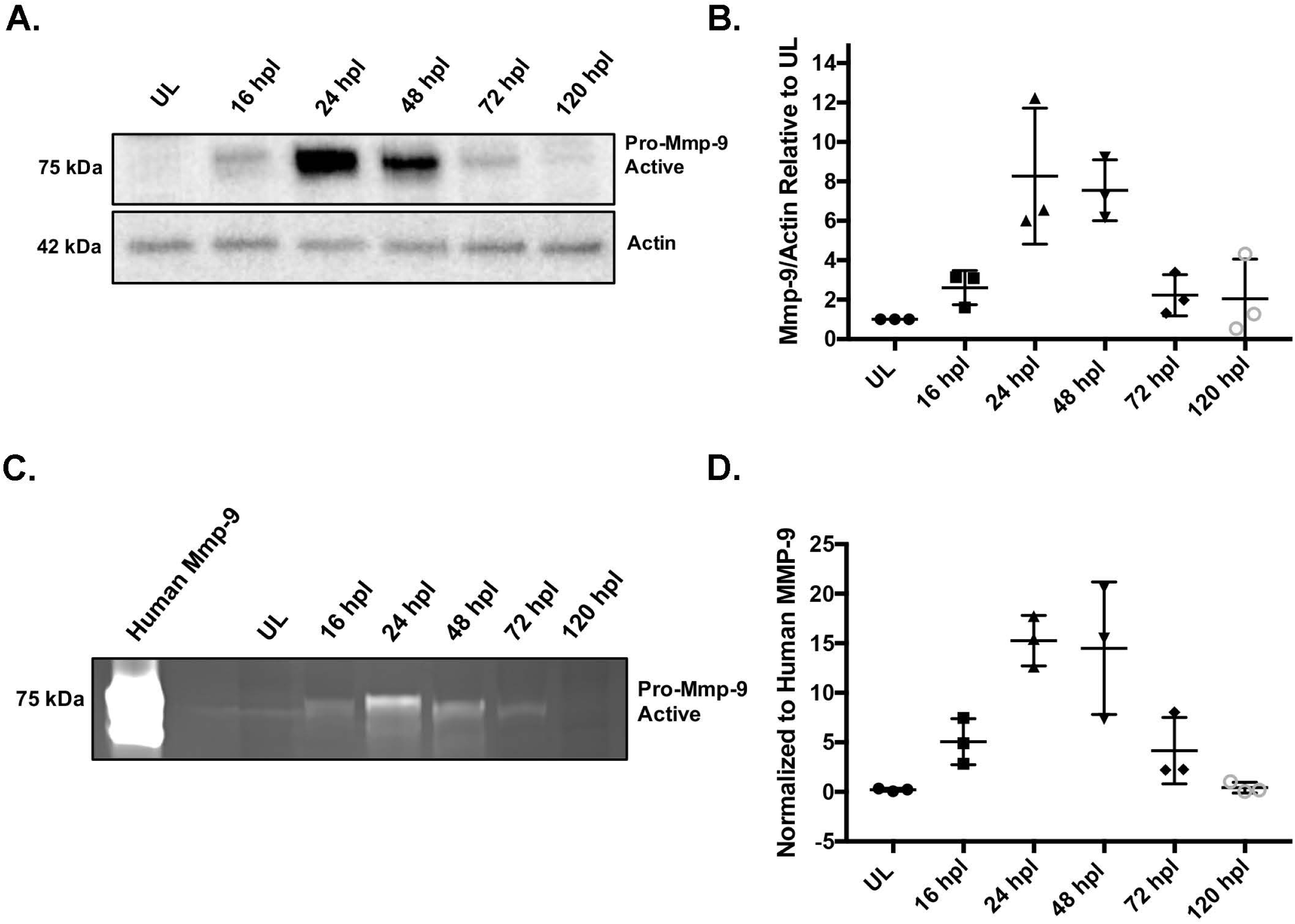
Mmp-9 is expressed and catalytically active following photoreceptor death. **(A)** Western blot of retinal proteins from unlesioned retinas (UL) and lesioned retinas at 16 hpl, 24 hpl, 48 hpl, 72 hpl, and 120 hpl, stained with anti-Mmp-9 and anti-actin (loading control) antibodies. **(B)** Densitometry of Mmp-9 protein, plotted relative to unlesioned controls. **(C)** Zymogram of Mmp-9 catalytic activity from unlesioned retinas (UL) and lesioned retinas at 16, 24, 48, 72 and 120hpl. Lanes in zymogram correspond to those in the Western blot. Purified human recombinant protein serves as a positive control. **(D)** Densitometry of Mmp-9 catalytic activity, plotted relative to the positive control.

Mmp-9 was previously named gelatinase B because its substrate, shared with Mmp-2, is denatured type-1 collagen (gelatin). In zymography, the polyacrylamide gel contains gelatin, and after protein renaturation and gelatin staining, areas of protease activity appear as clear, negatively-stained bands against the stained background (Chattopadhyay and Shubayev, 2009; Vandooren et al., 2013a). To evaluate the catalytic activity of Mmp-9, the same protein samples used for the Western blot analysis were used for zymography. Recombinant, active human MMP-9 was used as a positive control and shows strong catalytic activity as evidenced by the large negatively-stained band. Unlesioned retinas contain no detectable Mmp-9 catalytic activity. In contrast, the catalytic activity parallels the data from the Western blot analysis (ANOVA F-ratio = 11.870, p = .0003) (Figure 4C, D). Together, results from the Western blot and zymogram analyses show that catalytically-active Mmp-9 is induced by a lesion that selectively injures photoreceptors, and the time course of Mmp-9 synthesis and catalytic activity parallels the proliferative phases of photoreceptor regeneration.

### CRISPR mutants lack Mmp-9 protein and catalytic activity

To investigate the function of *mmp-9* during photoreceptor regeneration, mutants were generated using CRISPR-Cas9 (Hwang et al., 2013), targeted to the *mmp-9* catalytic domain in exon 2 (Figure 5A). The 19 base sgRNA produced several alleles, two of which were bred to homozygosity and characterized further, a 23 bp deletion (designated *mi5003*; ZFIN, https://zfin.org/) and an 8 bp insertion (designated *mi5004*; ZFIN, https://zfin.org/), respectively (Figure 5B). Individuals from the two lines were crossed to create compound heterozygotes to evaluate the potential for off target effects in either of the two independent lines (see next section). Each of the two *mmp-9* lines carries a mutation that results in a frameshift that predicts premature stop codons (Extended Data supporting Figure 5 labeled as Figure 5-1). Retinas from mutants were characterized by Western blot analysis and zymography. Unlesioned retinas had no detectable Mmp-9 or catalytic activity (Figure 5C, D). As expected, following photoreceptor ablation, Mmp-9 protein and catalytic activity are present in wild-type retinas (Figure 5C, D). In contrast, there was trace Mmp-9 and catalytic activity in the *mi5004* line, but no detectable Mmp-9 or catalytic activity in *mi5003* line (Figure 5C, D). Based on these results, subsequent regeneration experiments were conducting using the *mi5003* line, though the basic observations reported here were confirmed in the *mi5004* line (data not shown) and the compound heterozygotes (see below).

**Figure 5:**
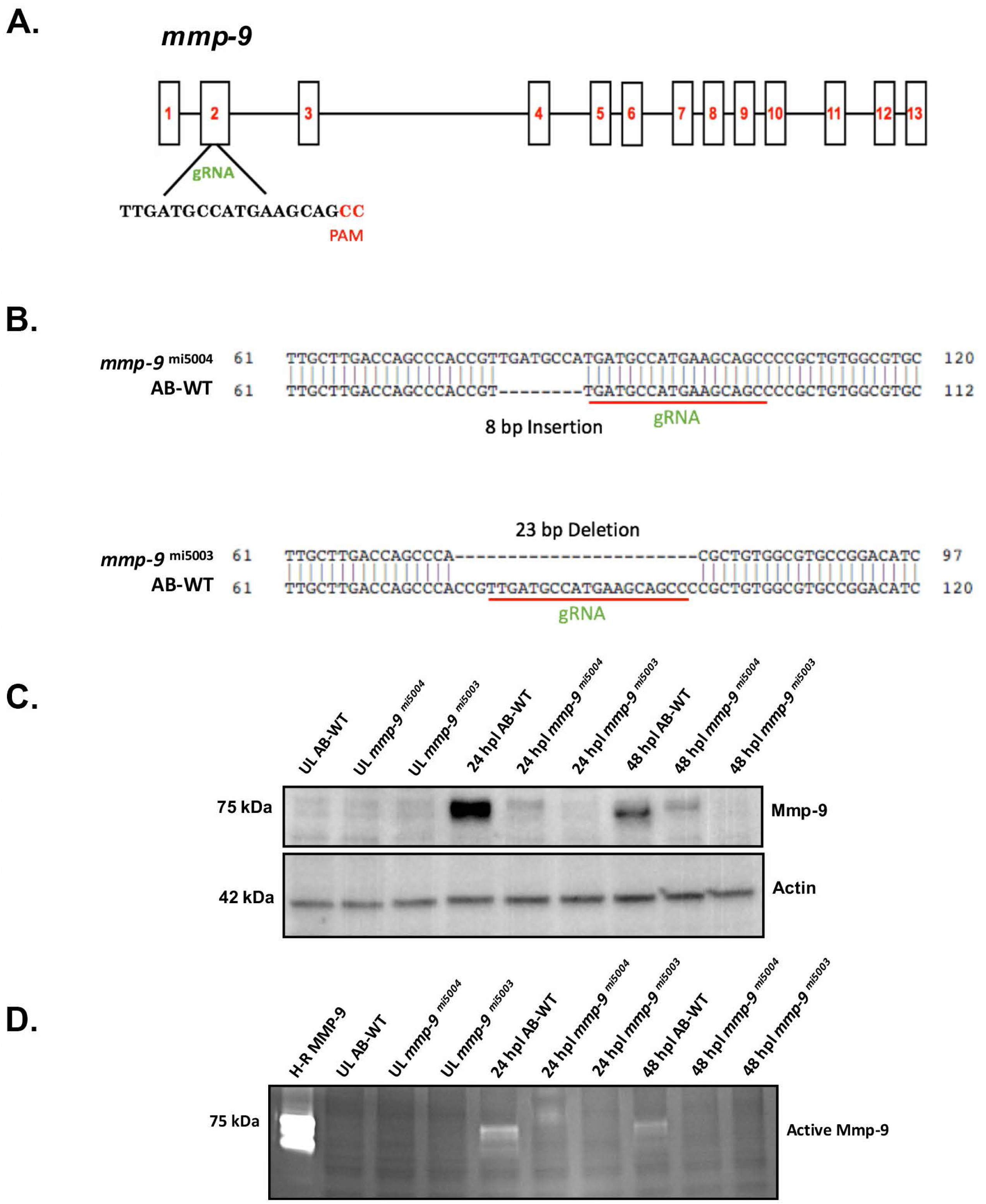
CRISPR mutants lack catalytically-active Mmp-9. **(A)** Genomic structure of *mmp-9* and the gRNA target sequence. **(B)** Sequence alignment for two indel mutations - 8 bp insertion (*mmp-9^mi5004^*) and 23 bp deletion (*mmp-9^mi5003^)*. Red underline indicates the sequence targeted by the 19 bp gRNA. **(C)** Western blot of retinal proteins from unlesioned retinas (UL wild-type; UL *mmp-9^mi5004^*; UL *mmp-9^mi5003^*) and lesioned retinas (wild-type; *mmp-9^mi5004^*; *mmp-9^mi5003^*) at 16, 24, and 48 hpl immunostained with anti-Mmp-9 and anti-actin (loading control) antibodies. **(D)** Zymogram of Mmp-9 catalytic activity. Lanes in zymogram correspond to those in the Western blot. Purified human recombinant protein serves as a positive control.

### In Mmp9 mutants, photoreceptor death results in overproduction of photoreceptor progenitors and regenerated photoreceptors

We first sought to determine if there were any differences in retinal development or adult starting points for the injury-induced proliferation in the *mmp-9* mutants. At 72 hpf, there were no differences between wildtype and the two mutant lines in the size of the eyes or degree of proliferation (Figure 6A, B, C), indicating early retinal development was not altered by the absence of Mmp9. Similarly, qualitative comparisons of adult retinas showed the appearance of wildtype and mutant retinas did not vary in any identifiable manner (Figure 7A). Finally, comparing the number of Müller glia and the number of mitotic progenitors, which support persistent neurogenesis in adult retinas (Raymond et al., 2006), showed there were no quantitative differences between wild-type animals and the *mi5003* line in the number of Müller glia or the level of intrinsic cell proliferation (Figure 7B,C).

**Figure 6.**
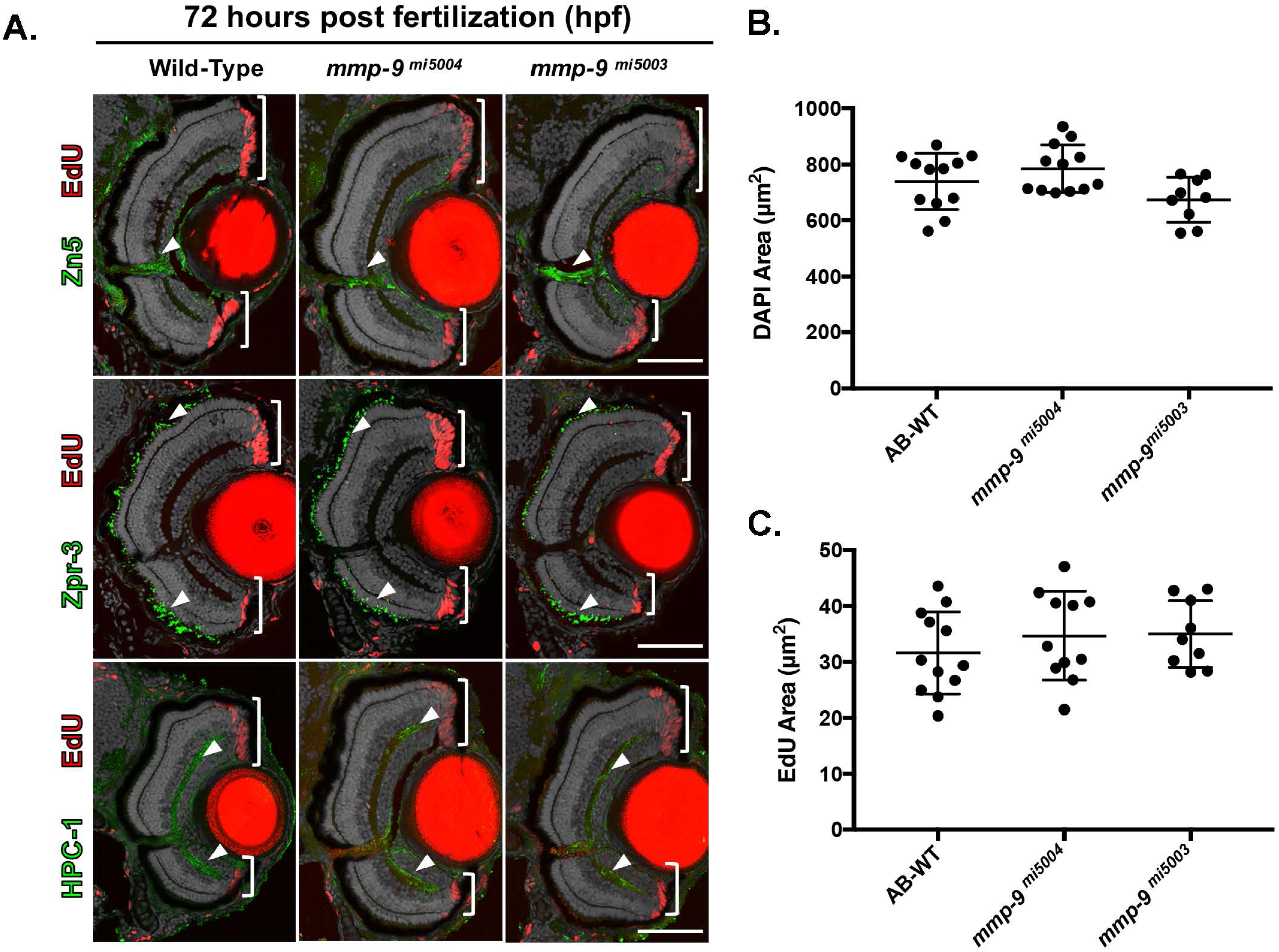
Retinal development is unaltered by the absence of mmp-9. **(A)** Sections from 72 hpf larvae immunostained (arrowheads) for ganglion cells (ZN5), rod photoreceptors (ZPR-3), and amacrine cells (HPC1). **(B)** Quantification of retinal area in wild-type (722.36 ±105.39 μm; n=12) compared to mmp9 mutants (676.95 ± 75.05 μm; n=9), p=.286. **(C)** Quantification of the area of EdU-labeled cells, as an indicator of proliferation, in wild-type (30 ± 7.84μm; n=12) and to mmp9 mutants (33.5 ± 6.59 μm; n=9), p=.293. Scale bar equals 25 ***μ***m.

**Figure 7.**
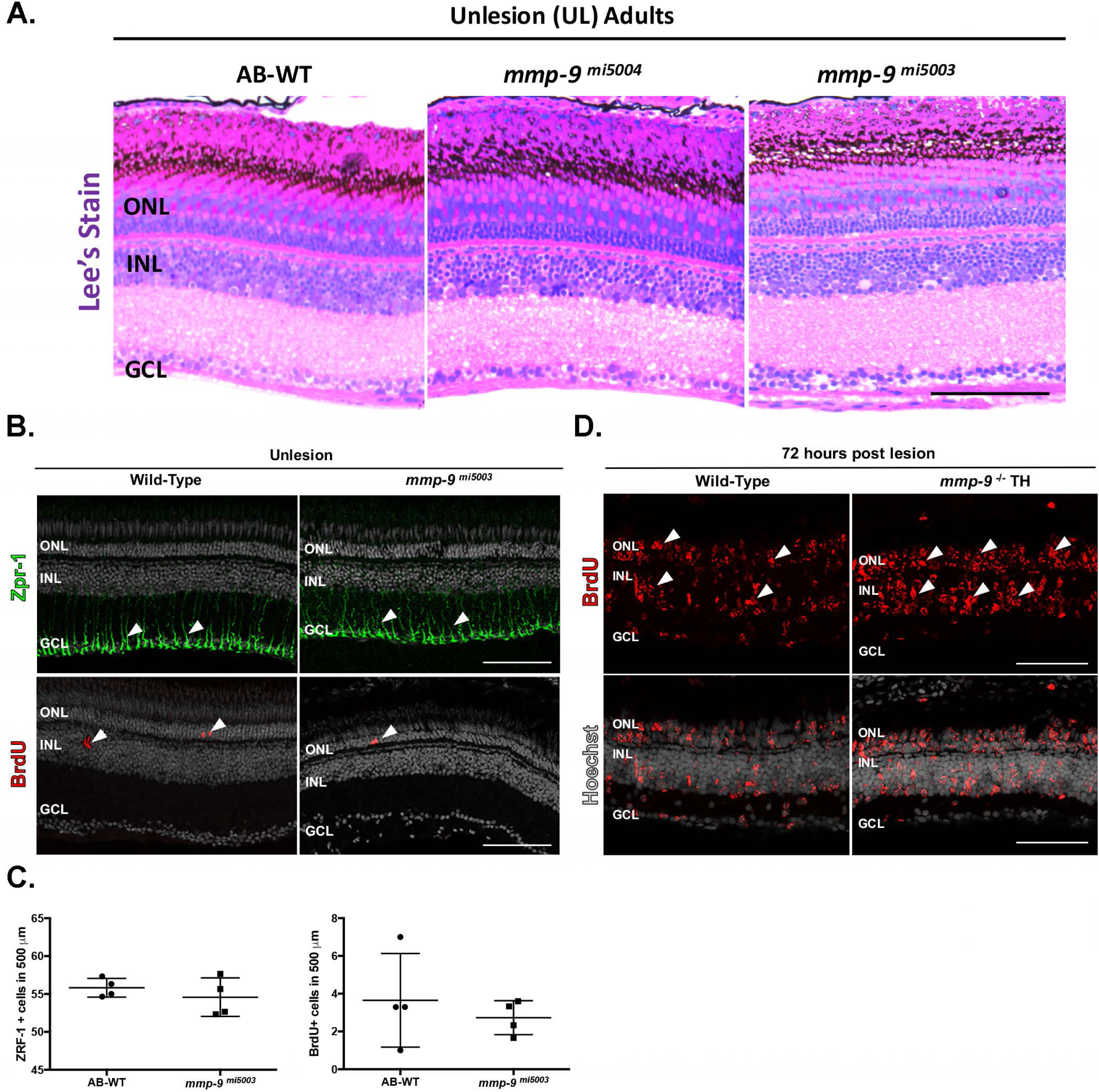
Anatomy of adult retina in Mmp-9 mutants. **(A)** Plastic sections illustrating cellular and synaptic layers in adult retinas from wildtype and Mmp-9 mutants. These slightly oblique sections also reveal elements of the cone photoreceptor mosaic. **(B)** Processes of Müller glia (green) stained with the zrf-1 antibody (top). BrdU-labeled cells (red, indicated by arrowheads) in wild-type and mutant retinas (bottom). **(B)** Immunostained Müller glia (top) and Brdu-labeled cells (bottom) in wild-type and mutant retinas. **(C)** Number of Müller glia in wild-type (55.83 ± 1.23; n=4) and mutant retinas (54.58 ± 2.54; n=4), p=.409 and dividing cells (bottom) in wild-type (3.66 ± 2.47; n=4) and mutant retinas (2.75 ± .92; n=4), p=.515. **(D)** BrdU-labeled cells (red) in wild-type (left) and transheterozygote (TH) retinas, qualitatively illustrating the overproduction of injury-induced progenitors in transheterozygotes. Scale bars equal 25***μ***m.

To determine the consequence of Mmp-9 loss-of-function on retinal regeneration, wild-type and mutant animals received photolytic lesions, Müller glia-derived progenitors were labeled with BrdU between 48-72hpl, and BrdU-labeled cells were counted at 72 hpl. Relative to wild-type animals, mutants had significantly more BrdU-labeled progenitors at 72 hpl (p = .0186) (Figure 8A, B). This over production of the Müller glia-derived progenitors was observed in both mutant lines and compound heterozygotes (Figure 7C), indicating that the hyperproliferation observed here is unlikely to be a consequence of off-target effects induced by the CRISPR-cas9 technique.

**Figure 8:**
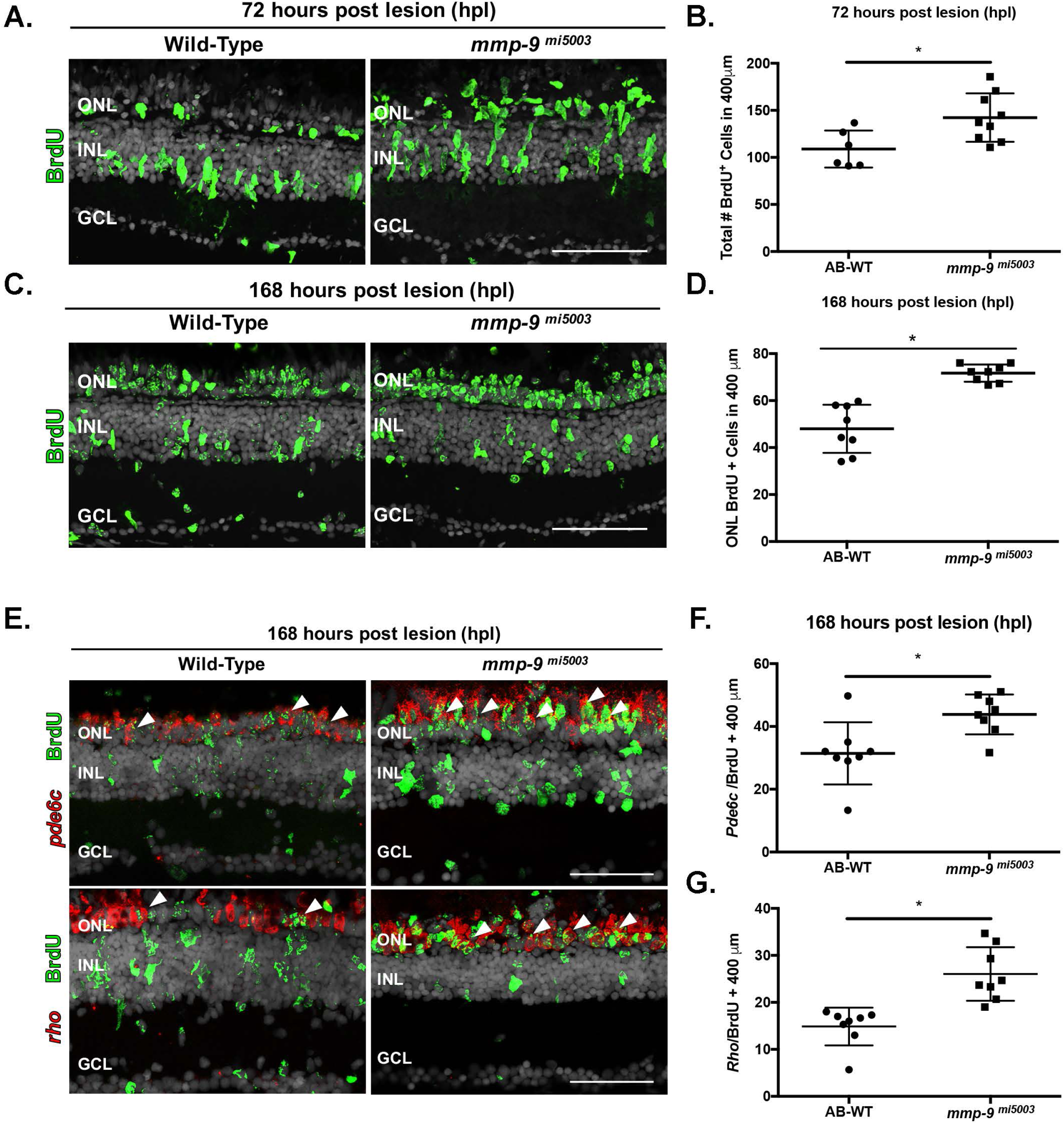
Photolytic lesions in *mmp-9* mutants results in the over production of injury-induced progenitors and regenerated photoreceptors. **(A)** BrdU-labeled cells (green) in wild-type and *mmp-9 ^mi5003^*. **(B)** Number of BrdU+ cells from wild-type (109 ± 19.66 cells; n=6) mutant retinas (142.3 ± 25.72 cells; n=9) at 72 hpl. *p = 0.0186. **(C)** BrdU-labeled cells (green) in wild-type and mutant retinas at 168 hpl. **(D)** Number of BrdU+ cells in the ONL of wild-type (48 ± 10.24 cells; n=8) and mutant retinas (71.71 ± 3.7 cells; n=8). *p=0.0001. **(E)** Double labeled, regenerated photoreceptors using *in situ* hybridization for rods (*rho) and cones (pde6c;* red signal) and BrdU (green) at 168hpl. **(F)** Number of regenerated cone photoreceptors in wild-type (31.42 ± 9.88 cell; n=8) and mutant retinas (43.83 ± 10.68 cells; n=8) at 168 hpl. *p=0.0301. **(G)** Number of regenerated rod photoreceptors in wild-type (14.88 ± 4.02 cells; n=8) and mutant retinas (26.04 ± 5.69 cells; n=8) at 168 hpl. *p=0.0005. ONL-outer nuclear layer; INL-inner nuclear layer; GCL-ganglion cell layer. Scale bars equal 50***μ***m.

It is well established that Mmp-9 plays a role in tissue remodeling and cell migration (Parks et al., 2004). No overt tissue remodeling was observed following photoreceptor death and during photoreceptor regeneration. To determine if Mmp-9 loss-of-function alters the migration of Müller glia-derived progenitors from the inner nuclear layer, where Müller glia reside, to the outer nuclear layer, which contains the photoreceptor nuclei, animals were subjected to photolytic lesions, exposed to BrdU between 48 and 72hpl and allowed to survive to 7dpl, a time point where proliferation and migration are complete and photoreceptor differentiation has commenced. Qualitative inspection shows that Mmp-9 loss-of-function does not alter the migration of the Müller glia-derived progenitors (Figure 8C). Further, cell counts show that the over production of photoreceptor progenitors observed in mutants at 72hpl results in significantly more BrdU-labeled cells within the ONL at 7dpl (p = .0001; Figure 9B, D). Further, at 7 dpl, there are significantly more regenerated rod and cone photoreceptors (p = .0301 and p = .0005, respectively; Figure 8E-G). Together, these data show that, in the absence of an injury, Mmp-9 loss-of-function has no impact retinal development, the proliferative potential of the adult retina or, following photoreceptor death, the migration of Müller glia-derived progenitors. The hyperproliferation of injury-induced progenitors in mutants indicates that Mmp-9 functions as a negative regulator of proliferation among Müller glia-derived progenitors, though the mechanistic details through which this is accomplished are not yet known (see Discussion). Further, the parallel overproduction of photoreceptors at 7dpl indicates that Mmp-9 plays no apparent role in governing the timing of cell cycle exit or the onset of photoreceptor differentiation.

**Fig 9.**
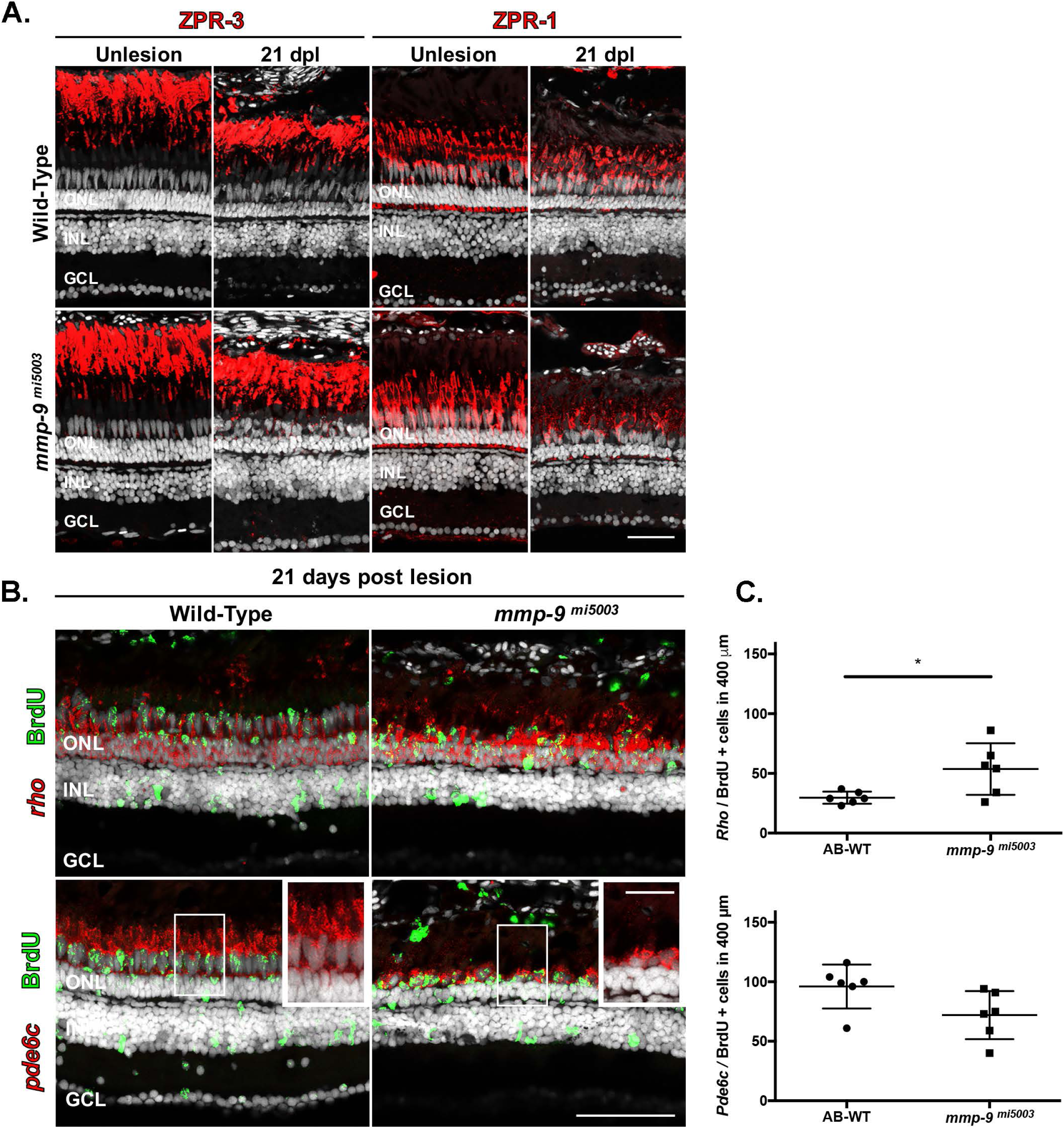
Mmp-9 regulates the maturation of regenerated photoreceptors. **(A)** Immunostaining for red-green double cones with ZPR-1 and rods with ZPR-3. Wild-type are in the top row; mutants are in the bottom row. Unlesioned retinas are left; lesioned retinas at21 dpl are right. **(B)** Double labeled, regenerated photoreceptors at 21dpl using *in situ* hybridization for rods (*rho) and cones (pde6c;* red signal) and BrdU (green). Insets detail the morphology of regenerated photoreceptors in wild-type (left) and mutant retinas (right). **(C)** Number of regenerated cone photoreceptors (bottom) in wild-type (95.81 ± 18.49 cells; n=6) and mutant retinas (71.89 ± 20.37 cells; n=6) at 21 dpl. p=0.059. Number of of regenerated rod photoreceptors (top) in wild-type (29.6 ± 4.94 cells; n=6) and mutant retinas (53.39 ± 21.62 cells; n=6) at 21 dpl. *p=0.0270. ONL-outer nuclear layer; INL-inner nuclear layer; GCL-ganglion cell layer. Scale bar equals 50***μ***m.

### Mmp-9 governs maturation and survival of regenerated cone photoreceptors

The qPCR data showed that *mmp-9* levels remain significantly elevated at the time injury-induced photoreceptor progenitors exit the cell cycle and begin differentiating into mature photoreceptors (Figure 1A). We infer that, though not detected by Western blot analysis, Mmp-9 protein persists as well. This suggests that Mmp-9 may have functional roles beyond regulating proliferation during the initial stages of photoreceptor regeneration. Therefore, regenerated rod and cone photoreceptors were qualitatively and quantitatively compared in wild-type and mutants at 21dpl, a time point where regeneration is complete (Powell et al., 2016). For both groups, retinal sections were labeled with rod- and cone-specific antibodies and regenerated photoreceptors were counted in sections and whole mount preparations. These analyses showed there were no qualitative differences between wild-type and mutants in the appearance of regenerated rod photoreceptors (Figure 9A), and the over production of rod photoreceptors observed at 7dpl was present at 21dpl (Figure 9B, C).

In mutant retinas at 21dpl, the maturation and survival of cones were clearly compromised. Relative to controls, regenerated cones in mutants have shorter outer segments and appear to be fewer in number (Figure 9A, insets 9B,C). Counts of regenerated cones in tissues sections show the initial over production of cones, evident at 7 dpl, is absent at 21 dpl (Figure 9C). Cones were also counted in retinal wholemounts (Figure 10). In both wild-type and mutant retinas, cone photoreceptors in unlesioned retinas are characterized by a very precise lattice mosaic, a feature of cone photoreceptors in teleost fish (Nagashima et al., 2017; Figure 10A). The number of cone photoreceptors in unlesioned retinas from wild type and the mutant retinas were invariant (Figure 10B). Following photoreceptor ablation and regeneration, areas of the retina that contain regenerated photoreceptors are identifiable by the marked spatial degradation of the mosaic, though individual cone photoreceptors remain readily identifiable (Nagashima et al., 2017). Counts of regenerated cones in wholemounts show that mutants have significantly fewer regenerated cones than wildtype animals (p=.0229) (Figure 10A, B). Finally, as an independent measure of the maturation of cone photoreceptors, Western blot analysis was performed using an antibody against the cone-specific transducin protein, Gnat-2 (Figure 10C; Lagman et al., 2015). As suggested by the invariant number of cones in unlesioned retinas, Gnat-2 levels in unlesioned retinas are comparable in wild-type and mutant animals. In lesioned retinas, Gnat-2 levels begin to recover in wild-type animals by 14 dpl, and by 21 dpl values in wild-type animals are nearly at the levels found in unlesioned retinas (Figure 10C, D). In contrast, Gnat-2 levels in mutants lag behind wild-type levels, and at 21dpl Gnat-2 levels in mutant animals are significantly less than in wild-types (p = .0014) (Figure 10C, D). Comparable measures specific to rod photoreceptors showed there were no differences between wild-type and mutant animals (data not shown). These results indicate that Mmp-9 also plays a functional role during photoreceptor regeneration, well after injury-induced proliferation and the initial differentiation of regenerated photoreceptors is complete. Further, this function is specific to cones.

**Fig 10.**
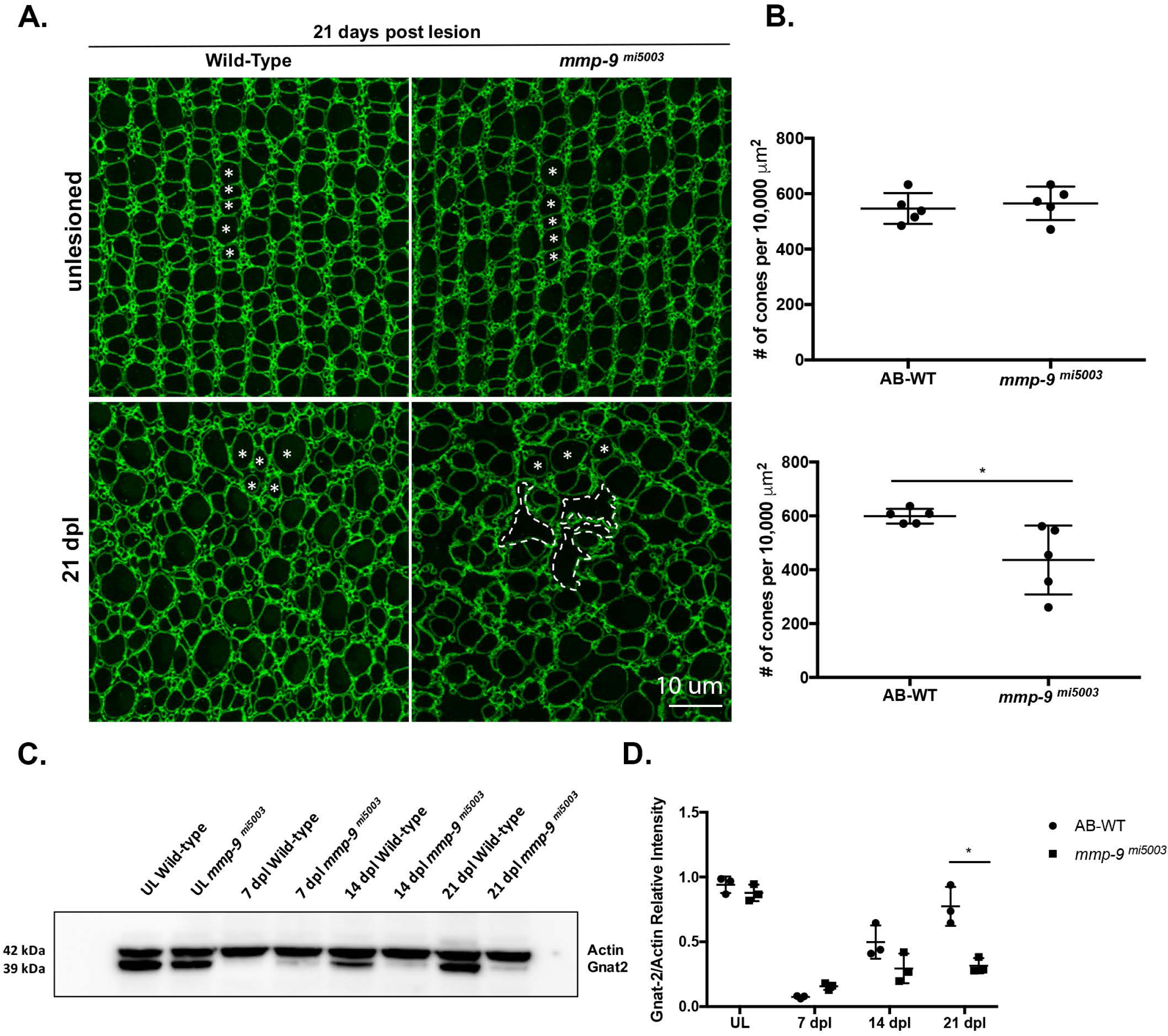
Mmp-9 is required for the survival of regenerated cone photoreceptors. **(A)** Wholemounts of wild-type (left) and mutant retinas (right) immunostained for ZO-1. Unlesioned retinas are top; lesioned retinas at 21dpl are bottom. Asterisks indicate profiles of cones. In mutants, gaps due to missing cones are replaced by the irregular apical processes of Müller glia (dashed lines). **(B)** Number of unlesioned cones from wild-type (546.49 ± 55.69; n=5) and *mmp-9 ^mi5003^* (565.33 ± 27.42; n=5) *p=.516; below the number of regenerated cones from wild-type (599.11 ± 27.42 cones; n=5) and mutant retinas (436.09 ± 128.04 cones; n=5) at 21 dpl. *p=0.0238. **(C)** Western blot of retinas stained with antibodies against gnat-2 and actin at 7, 14, and 21 dpl. **(D)** Densitometry of gnat-2 labeling in the Western blot. A significant difference in gnat-2 levels were observed at 21 dpl. *p=.0014. Scale bar equals 10 ***μ***m.

### Late anti-inflammatory treatment rescues the maturation and survival defect of regenerated cones in *mmp-9* mutants

MMP-9 is known to cleave inflammatory cytokines, and we hypothesize it may function to modify cytokine signaling during the resolution phase of tissue inflammation. To determine if the defects in the maturation and survival of cone photoreceptors is a consequence of persistent inflammation, we treated mutants with vehicle or Dex between 3 and 13dpl to suppress the inflammatory response, beginning after the initial formation of Müller glia-derived progenitors and throughout the phase of photoreceptor differentiation and maturation (Figure 11A). Cone photoreceptors were then counted in retinal wholemounts at 21dpl from vehicle and Dex-treated mutants. The number of cones at 21dpl in vehicle-treated mutants approximates that shown above (Figure 10B; Figure 11mB, C). In contrast, there were significantly more regenerated cones in the Dex-treated mutants than in the vehicle-treated controls (Figure 11B, C; p =.01), restoring the number of regenerated cones nearly to that observed in wild-type animals. Further, regenerated cones in the Dex-treated mutants were noticeably more mature than in controls, as evidenced by their more regular arrangement and increased overall length (Figure 11D). Correlated with these data, activated microglia are present in the ONL of vehicle-treated mutants (Figure 11E), a notable feature of the early stages of photoreceptor death and a hallmark of inflammation in the retina (White et al., 2017). These results suggest that the absence of Mmp-9 results in a persistent immune activation, which selectively compromises the survival and maturation of cone photoreceptors.

**Fig 11.**
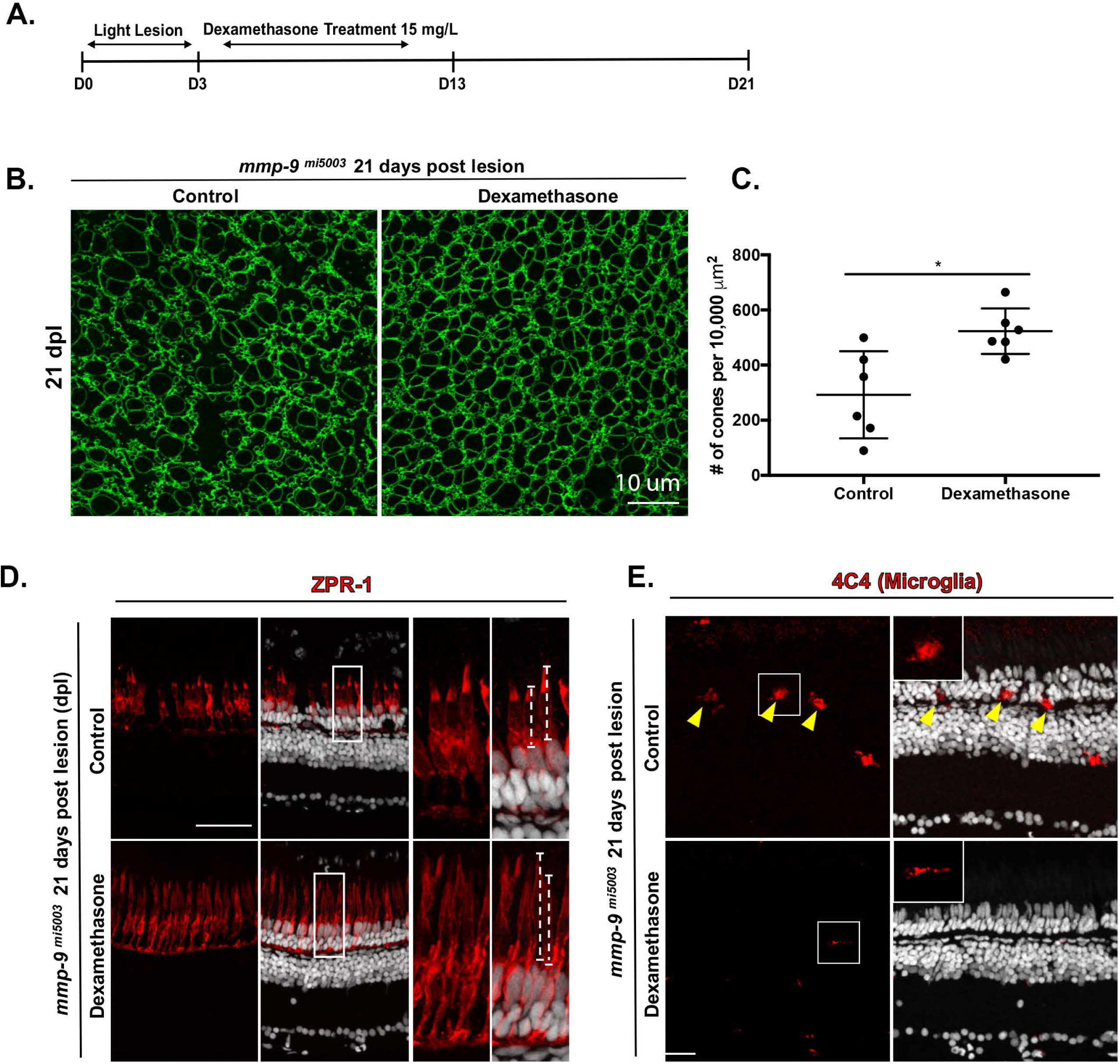
Anti-inflammatory treatment rescues the maturation and number of regenerated cones in mutants at 21dpl. **(A)** Experimental paradigm for the photolytic lesions and Dexamethasone treatment. **(B)** Wholemounts of mutant retinas immunostained for ZO-1. Control retina is left; Dex-treated retina is right. **(C)** Number of regenerated cones in control (292.20 ± 158.11 cones; n=6) and Dex-treated retinas (522.79 ± 82.55 cones; n=6) at 21 dpl. *p=0.0010. **(D)** Immunostaining with ZPR-1 for red-green double cones in control (top) and Dex mutants (bottom) 21 dpl. Insets illustrate differences in the maturation and lengths of the cone photoreceptors (dashed lines; control, 22.1 ± 7.2μm, n=40 cells; Dex-treated, 29.7 ± 4.8μm, n=69 cells; p<.001) **(E)** Immunostaining for microglia using the 4C4 antibody in control (top) and Dex-treated retinas (bottom). Inserts illustrate ameboid (top) and ramified (bottom) microglia. Scale bars equal 10***μ***m in panel A and 40***μ***m in panels D and E.

**Fig 12.**
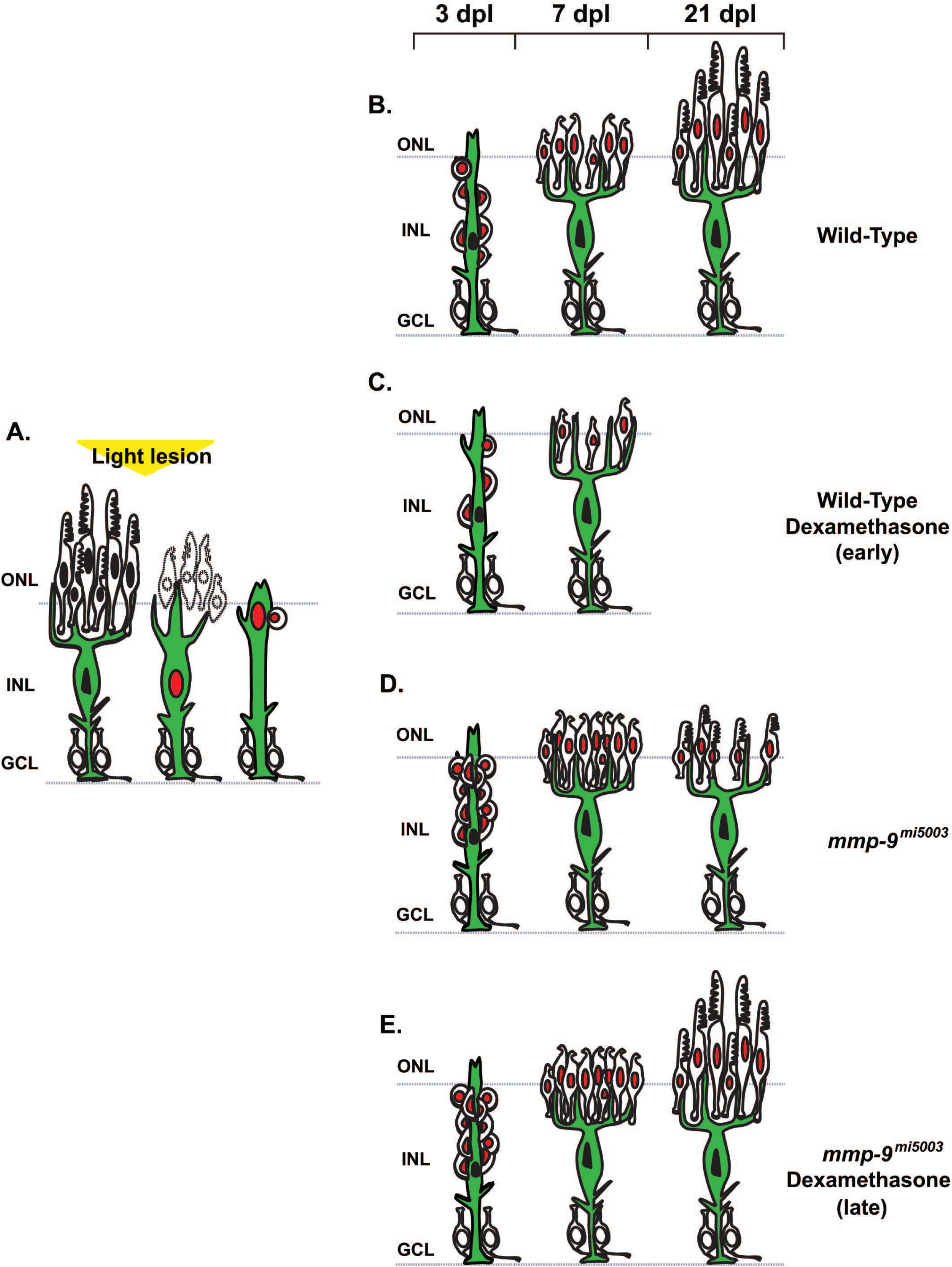
Summary Diagram of cone regeneration. **(A)** Müller glia respond to photoreceptor death by expressing *mmp-9*, undergoing interkinetic nuclear migration and a single asymmetric cell division that gives rise to a neuronal progenitor. **(B)** In wild-type animals, neuronal progenitors form a neurogenic cluster around the Müller glia (3 dpl), migrate to the ONL, and differentiate into cone photoreceptors (7 dpl) that then mature (21 dpl). **(C)** Anti-inflammatory treatment results in fewer Müller gli-derived progenitors and fewer regenerated photoreceptors. **(D)** In the absence of Mmp-9, there is overproduction of Müller glia-derived progenitors and regenerating photoreceptors. However, at 21dpl, cone maturation and survival are compromised. **(E)** In the absence of Mmp-9, anti-inflammatory treatment rescues the maturation and survival of cone photoreceptors. ONL-outer nuclear layer; INL-inner nuclear layer; GCL-ganglion cell layer.

## DISCUSSION

The results of this study are summarized in Figure 13. In response to photoreceptor cell death, Müller glia undergo a single asymmetric division through interkinetic nuclear migration to produce progenitors, which rapidly proliferate, migrate and differentiate to replace the ablated photoreceptors (Figure 13A, B). Dexamethasone-induced immunosuppression inhibits proliferation, resulting in diminished photoreceptor regeneration (Figure 13C). Loss of Mmp-9 function effectively reverses the consequences of immunosuppression, resulting in hyperproliferation and overproduction of regenerated photoreceptors (Figure 13D). However, the absence of Mmp-9 leads to defects in the survival and maturation of cone photoreceptors (Figure 13D). Immunosuppression in mutants following photoreceptor regeneration rescues the maturation and survival defects (Figure 13E). Based on these results, we conclude that inflammation is required during the proliferative phase of photoreceptor regeneration, and as a component of the inflammatory response, Mmp-9 provides an inhibitory balance that refines this proliferative response. Finally, we conclude that Mmp-9 is required for the survival and maturation of regenerated cone photoreceptors.

Immune system activation is required for regenerative neurogenesis (Fischer et al., 2014; Kyritsis et al., 2012; White et al., 2017; Caldwell et al., 2019). In the forebrain and brainstem of adult zebrafish, immunosuppression inhibits the production of progenitors after traumatic lesion (Kyritsis et al., 2012; Caldwell et al., 2019). In the retina of larval zebrafish, immunosuppression results in decreased migration of microglia into the outer retina during the regeneration of rod photoreceptors (White et al., 2017). In the chick, ablation of microglia completely suppresses the formation of Müller glia-derived retinal progenitors (Fischer et al., 2014). Our data are consistent with these reports and indicate that acute inflammation is a required component of neuronal regeneration in the vertebrate central nervous system.

Based on their structure and specific substrates, MMPs are classified into six groups: gelatinases, collagenases, MTMMPs, stromelysins, and matrilysins (Vandooren, et al., 2013b). MMP-9 is a secreted protease that is a member of the gelatinases. In response to photoreceptor injury and death, *mmp-9* is rapidly induced in Müller glia, and anti-inflammatory treatment significantly suppresses *mmp-9* expression. These data suggest Mmp-9 as a component of the inflammatory response in the retina. Several previous reports showed that following neuronal death Müller glia secrete the inflammatory cytokines, *tnf-α*, *leptin*, and *il-6* (Nelson et al., 2013; Zhao et al., 2014). Recent transcriptome and gene ontology analysis for Müller glia, isolated during photoreceptor injury and death, identified cytokine signaling and the immune response as activated pathways (Roesch et al., 2012; Sifuentes et al., 2016). However, in the mammalian retina, neuronal death leads to persistent active gliosis in Müller glia and failure of the spontaneous expression of intrinsic reprogramming factors. This limits the ability for neuronal regeneration (Bringmann et al., 2006, 2009; Reichenbach and Bringmann, 2013; see also Ueki et al., 2015). Our data add details to the innate immune response in the retina that is required for stem cell-based regeneration of photoreceptors. It is interesting to note that the activation of cytokine pathways is conserved in the transcriptomes of Müller glia in both zebrafish and mouse models of photoreceptor degeneration (Roesch et al., 2012; Sifuentes et al., 2016), suggesting that photoreceptor death in vertebrates results in a common inflammatory response among Müller glia.

Mmp-9 loss-of-function results in an increased number of Müller glia-derived progenitors, resulting initially in more regenerated photoreceptors. The overproduction of regenerated photoreceptors is interpreted simply to result from the overproduction of progenitors, and Mmp-9 plays no role in the timing of cell cycle exit or initial photoreceptor differentiation. There are at least three interpretations that explain the overproduction of photoreceptor progenitors. First, the absence of Mmp-9 may allow more Müller glia to enter the cell cycle. Previous studies showed that neuronal damage results in only 65-75% of the Müller glia entering the cell cycle (Nagashima et al., 2013). However, we demonstrated that *mmp-9* was expressed only by dividing Müller glia. The absence of proliferation in *mmp-9*-negative Müller glia argues against the possibility that additional Müller glia are recruited to the cell cycle. Second, in the *mmp-9* mutants Müller glia may undergo more than a one round of cell division, thereby increasing the size of the initial pool of Müller glia-derived progenitors, which then divide at a normal rate. We do not favor this possibility, given a single asymmetric division among Müller glia is a hallmark of regenerative neurogenesis in the zebrafish retina. Third, in the *mmp-9* mutants, proliferation of the Müller glia-derived progenitors is accelerated. We view this the likeliest of the three possibilities, and hypothesize that, as an extracellular protease, Mmp-9 regulates the concentration of cytokines and/or growth factors that regulate cell cycle kinetics within the niche of progenitors that cluster around each parental Müller glia, and in the absence of Mmp-9, mitogenic signals persist, leading to accelerated proliferation (Bosak et al., 2018; Le et al., 2007; Manicone and Mcguire, 2008; Parks et al., 2004; Luo et al., 2012; Vandooren et al., 2013b). Finally, we speculate that in regenerating tissues the magnitude of the inflammatory response serves to quantitatively match the number of injury-induced progenitors to the degree of the cell death.

White et al, (2017) showed that in larval zebrafish the timing of immune suppression can delay or accelerate the regeneration of rod photoreceptors. As a pre-treatment, dexamethasone suppresses the regeneration of rods. This result aligns with what we report here for adults, inflammatory events are required for the activation of Müller glia and Müller glia-derived progenitors. However, if dexamethasone treatment commences after rod photoreceptor ablation (White et al., 2017), regeneration kinetics are enhanced. The enhanced regeneration following delayed immune suppression parallels the results from the Mmp-9 mutants, but it is unlikely a common mechanism can readily account for both results. The molecular milieu is complex in injured tissues, and the potential parallels between delayed suppression of inflammation and the absence of Mmp-9 await further characterization.

Recently, Kaur et al (2018) demonstrated a regulatory feedback loop involving Shh signaling and Mmp-9 during retinal regeneration in zebrafish. Pharmacological and genetic suppression of Shh signaling inhibited the proliferative response of Müller glia and their progenitors. This was accompanied by up-regulation of *mmp-9* and the repressor of proliferation, *insm1a* (Kaur et al., 2018; Ramachandran et al., 2012). The hyperproliferation we observed in the *mmp-9* mutants is consistent with their proposed model. When Shh signaling declines, progenitors increase secreted Mmp-9, which then degrades extracellular factors that facilitate proliferative competence, resulting in increased *insm1a* and exit from cell cycle. In this model, the absence of Mmp-9 would fail to degrade factors that stimulate proliferation, leading to hyperproliferation. The positive feedback loop between Shh signaling and Mmp-9 suggests that Shh and HB-EGB, factors that function upstream of *insm1a*, may be proteolytic substrates for the Mmp-9 (Wan et al., 2012).

An inconsistency in the Kauer et al., (2018) study is their observation that morpholino inhibition of Mmp-9 action decreases the production of Müller glia-derived progenitors. This report is inconsistent with our data from the *mmp-9* mutants and raises the possibility that (1) the reliance on morpholinos, with the recognized potential for off-target effects (Schulte-Merker and Stainier, 2014; Kok et al., 2015; Stanier et al., 2017), may be the origin of these inconsistent data, or (2) their injury paradigm, which results in bloodborne factors and cells (White et al., 2017) invading the retina, may result in the activation of signaling pathways not present in the retina following a photolytic lesion.

In the Mmp-9 loss-of-function mutants, survival and maturation of regenerated cone photoreceptors are compromised. It is well established that chronic inflammation is a risk factor for photoreceptor dystrophies in humans, such as retinitis pigmentosa and age-related macular degeneration (AMD; Kauppinen et al., 2016; Chen and Xu, 2012; Whitcup et al., 2013; Yoshida et al., 2013). A subset of patients with AMD exhibit single nucleotide polymorphism (SNP) variants in the *MMP-9* gene, and elevated inflammation is associated with AMD pathogenesis (Fritsche et al., 2016; Kauppinen et al., 2016; Parks et al., 2004). Mutational variants in the inflammatory regulators, TIMP-3, TGFB, TNF-α are also identified in human patients with AMD (Fritsche et al., 2016). We hypothesize that during photoreceptor regeneration in zebrafish, Mmp-9 functions late to resolve the acute inflammation stimulated by death of the photoreceptors, and, in the *mmp-9* mutants, there is prolonged inflammation that is selectively damaging to cones. The *mmp-9* mutants studied here may recapitulate elements of the pathologies that are characteristic of inflammatory photoreceptor diseases in humans, including AMD. Our study begins to shed light on the functional roles of Mmp-9 in the retina and a potentially important link between intrinsic retinal immunity, human photoreceptor dystrophies and photoreceptor regeneration.

## Acknowledgements

This work was supported by grants from the National Institutes of Health - R01EY07060 (PFH), R01 EY024519 (DRH), T32EY013934 (NJS), P30EYO7003 (PFH) and an unrestricted grant from the Research to Prevent Blindness, New York. The authors thank Dilip Pawar for technical assistance. Fish lines and reagents provided by ZIRC were supported by NIH-NCRR Grant P40 RR01.

---

**Figure 5-1.**
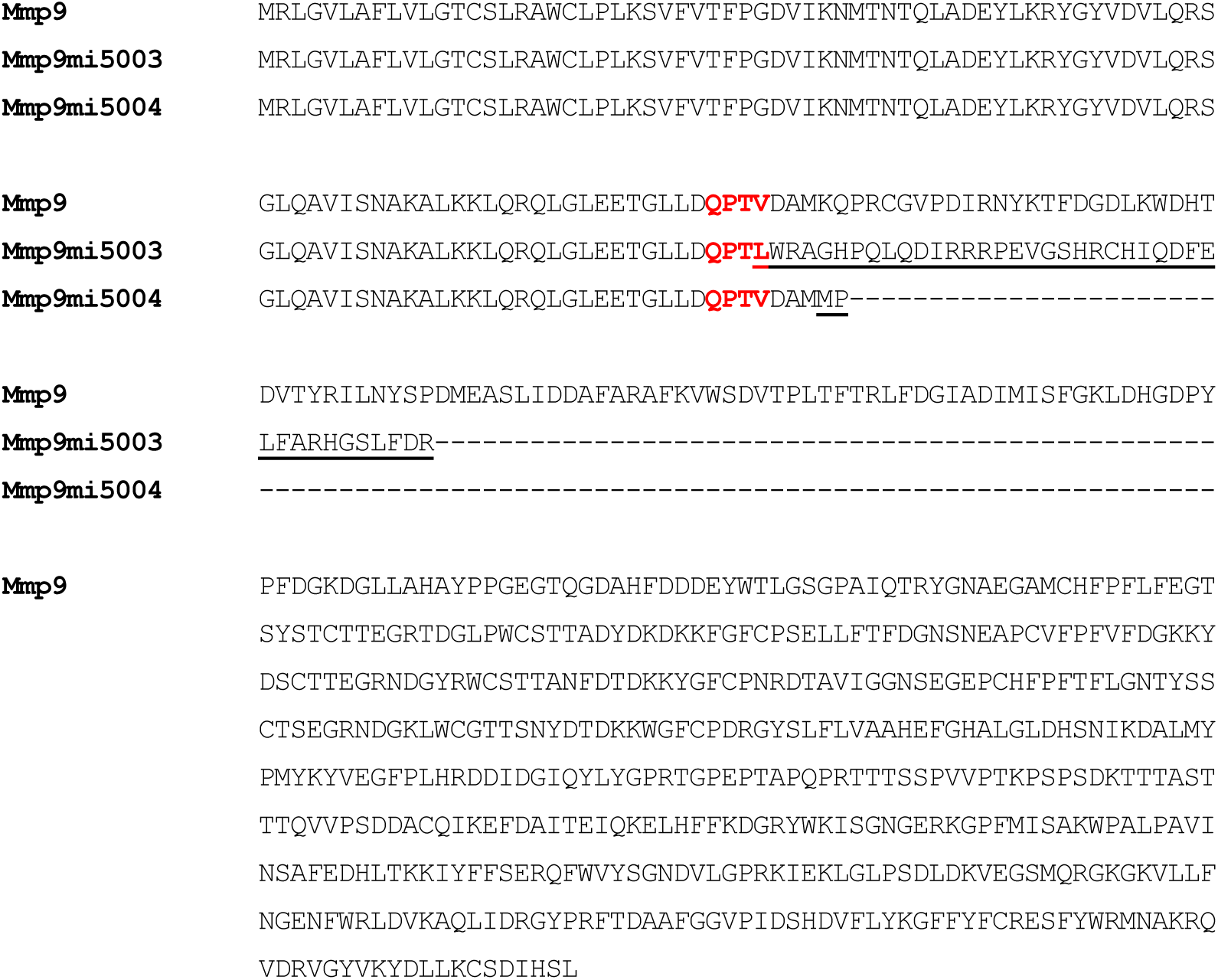
Extended Data supporting Figure 5. **(A)** Amino acid sequence alignments for Mmp-9 in wild-type and *mmp-9^mi5003^* and *mmp-9^mi5004^* mutants. The red amino acids denote the target of the gRNA. Insertion or deletion introduced a frameshift resulting in alteration of the amino acid sequence (*mmp-9^mi5003^)* or protein truncation (*mmp-9^mi5004^*).

